# Updated neuronal numbers of the rat hippocampal formation: redesigning the hippocampal model

**DOI:** 10.1101/2025.04.16.649153

**Authors:** Jon I Arellano, Pasko Rakic

**Affiliations:** Department of Neuroscience, Yale University, New Haven, CT, United States; Kavli Institute for Neuroscience at Yale, Yale University, New Haven, CT, United States

**Keywords:** hippocampus, neuron number, rat, stereology, entorhinal, subiculum

## Abstract

The hippocampal formation is a functional entity that includes the hippocampus, subicular complex and the entorhinal cortex, and has an essential role in learning and memory, emotional processing and spatial coding. The well-defined structure of hippocampal fields and the segregation of the connections have made this structure a favorite candidate for functional models, that rely on fundamental information such as the number of neurons populating the hippocampal fields. Existing models on the rat rely on neuronal populations obtained from single studies, so we aimed to obtain more representative estimates by analyzing all available data. We identified 89 studies using reliable methodology that provided 264 stereological estimates of principal neuron populations. The resulting averages for males showed 1,000,000 neurons for the granule cell layer (GCL); 50,000 for the hilus; 210,000 for CA3; ∼30,000 for CA2; 350,000 for CA1 and 300,000 for the Subiculum. Entorhinal cortex (EC) averages for both sexes showed 108,000 neurons in layer II; 270,000 in layer III and 340,000 in layer V/VI. Most of those estimates are significantly different from those traditionally used in hippocampal models (e.g.: ∼2-fold difference in EC layer II), revealing an updated architecture of the rat hippocampal formation that might help build more realistic models of hippocampal connectivity and function. Comparisons by age or sex were not reliable given the scarce data available from adolescents or females, while comparisons by strain showed inconsistent results, with similar populations in most fields but significant differences in CA3/CA2. The reliability of this finding is discussed.

## Introduction

The hippocampal formation plays a critical role in memory consolidation, emotional processing, and spatial navigation, making it a central subject of physiological and functional research. It comprises the hippocampus, subicular complex and entorhinal cortex, which are interconnected by strong, spatially segregated, mostly unidirectional and bilateral connections. These characteristics make it an ideal candidate for connectivity and functional models, which require precise quantitative data on neuronal populations and synaptic connectivity.

A seminal review by (Amaral et al., 1990) provided a foundational quantitative description of hippocampal structure and connectivity in the rat, based on the limited studies available at the time, serving as a reference for numerous models up to the present (Treves and Rolls, 1992, 1994; Patton and McNaughton, 1995; Henze et al., 2002; Aimone et al., 2006, 2009; Leutgeb et al., 2007; Kempermann, 2011; Krueppel et al., 2011; Snyder and Cameron, 2012; Rolls, 2013; Newman and Hasselmo, 2014; Cole et al., 2020; Berdugo-Vega et al., 2023; Borzello et al., 2023; Vandael and Jonas, 2024). However, numerous subsequent studies have expanded upon these estimates, providing a more comprehensive picture of neuronal populations within the hippocampal formation.

To this end, we performed a systematic review and found 166 studies describing the population of principal neurons in the fields of the hippocampal formation of the rat. Review of the methodology and results (see material and methods) led to a selection of 87 studies containing 264 stereological estimates of the neuronal population of the granular cell layer (GCL) and hilus of the dentate gyrus (DG), stratum pyramidale of CA3, CA2 CA1, the subicular complex including subiculum, pre-, para- and post-subiculum and layers II, III and V/VI of the medial and lateral entorhinal cortex (EC). The dataset predominantly includes data from males (∼80%) as females are typically avoided in hippocampal studies due to concerns over hormonal influences ((Gould et al., 1990; Zhang et al., 2008a). Also, there was a bias towards young animals (6 months or younger) which represent almost three quarters of the datapoints, reflecting a historical youth bias in rodent research (Snyder, 2019; Arellano et al., 2024). Regarding strains, about 83% of estimates originated from Wistar or Sprague-Dawley (SD) rats, while the rest come evenly from Fischer 344 (F344) and Long Evans (LE) rats. In summary, more than half of the estimates (54%) come from Wistar or SD males up to 6 months of age.

A consistent feature of the data is the large variability between neuron estimates, that orbits around 2-2.5-fold but can reach almost 3-fold. Another characteristic is the scarcity of female data, typically avoided in hippocampal studies due to concerns over hormonal influences (Gould et al., 1990; Zhang et al., 2008a). Only the GCL reached 5 female estimates with similar average to the males. And in the entorhinal cortex (EC), we found few (n=3-5) estimates of layer populations coming from both males and females, showing that females tend to show 10-15% less neurons than males. For the rest of the fields, averages are based on males, and sex dimorphism, a feature consistently described in the rodent hippocampus (Madeira et al., 1992; Andrade et al., 2000; Nuñez et al., 2003b; Smith et al., 2008), precluding reliable comparisons of neuronal populations by sex.

The average number of neurons in the GCL was 1,050,000 in adult males and 1,010,000 in females, with an average of 1,030,000 for both sexes. In most other fields, averages were calculated only for adult males, as females were scarce. Our results show 50,000 neurons for the hilus; 210,000 for CA3; ∼30,000 for CA2; 340,000 for CA1 and 300,000 for the Subiculum. In the EC, males exhibited 116,000 neurons in layer II, 290,000 in layer III and 363,000 in layer V/VI, while females had 102,000, 247,000 and 313,000 respectively, representing about 85% of the estimates in males. When the medial and lateral EC areas (MEC and LEC, respectively) are considered, almost all data originate from females and is surprisingly consistent. MEC layer II shows about 60,000 neurons, layer III has about 140,000 and layer V/VI about 180,000. LEC layer II has 40,000 neurons, while layer III shows 110,000 and layer V/VI about 130,000, indicating that LEC has about 75% of the neurons present in MEC.

While the updated average for the GCL is very similar to the traditional estimate, the updated averages for the rest of fields (hilus, CA fields, subiculum and EC) differ significantly from those used traditionally as described by (Amaral et al., 1990), with values that reach 2-fold difference (e.g.: EC layer II or the subiculum). Also, we estimated the separate populations of CA3 and CA2, a differentiation described by (Amaral et al., 1990) that has been lost in most subsequent studies, as both fields are typically described together, although they have been reported to exhibit important differences in connectivity and function.

Most of the selected studies (85%) used the optical fractionator (OF; (West et al., 1991)), considered the most robust method to quantify particles, as it does not require volume estimates that add additional sources of variance to the estimate (Slomianka, 2021). The remaining 15% used the Nv*Vref model, that requires calculating the density of neurons (Nv) and the volume of structure of interest (Vref). We found also 9 older studies using assumption-based stereology (ABS) that were excluded as its methodology is not as robust as in DBS studies. In addition, those older studies tended to use developing animals instead of adults, and frequently mixed males and females, that are not the focus of our analysis.

Comparisons between strains have shown heterogeneous results across studies (Boss et al., 1985, 1987; Seress, 1988). Our dataset allowed us to compare Wistar and SD strains, as they were the most used. Results showed differences within 5% in the hilus and CA1 that reached 9% in the GCL and a statistically significant 23% difference in CA3/CA2, although the reliability of that significant difference might be questionable as described in the results and discussion.

Several reports have shown that adolescent rats tend to exhibit less neurons than adults in CA fields (Madeira et al., 1992; Wakuda et al., 2008; Elibol-Can et al., 2014), while others show no clear differences (De Araujo Furtado et al., 2024). While such differences are expected in the dentate gyrus, where neurogenesis is still significant during adolescence and young adulthood (Arellano and Rakic, 2024), it is surprising in the rest of the hippocampal fields, as neurons are generated largely pre- and perinatally and by the onset of adolescence, around 3 weeks of age (Arellano et al., 2024), the number of cells identifiable as neurons is expected to match adult levels. Our dataset contained only scarce adolescent data that tend to show lower values than adults, but their limited numbers precluded reliable comparisons.

Overall, the data presented here represent the most comprehensive collection of neuron number estimates for the rat hippocampal formation to date, and we hope it will provide a valuable resource to update neuronal populations for hippocampal models and facilitate comparative studies across species.

## Methods

### Field segregation and nomenclature

The hippocampal formation consists of three main structures: the hippocampus, the subicular complex and the entorhinal cortex. The hippocampus contains the dentate gyrus (DG) and the hippocampus proper also known as Ammon’s horn or Cornu Ammonis (CA) fields CA1-CA3. The dentate gyrus has two cellular layers: the granular cell layer (GCL), composed of mostly granular cells, and the polymorphic layer or hilus, that consists of a narrow territory inside the dentate blades, between the GCL and CA3 (Maliković et al., 2022).

The demarcation of the territories of the CA fields requires some detailed explanations. Cajal described two regions in the Ammon’s horn: regio inferior, populated by large pyramidal cells near the DG, and regio superior, composed of smaller pyramidal cells proximal to the subiculum (Ramon y Cajal, 1893). Later on, (Lorente de Nó, 1934) described 4 subfields: CA4, CA3, CA2 and CA1. Field CA4 corresponded to the pyramidal cells located inside the DG blades, (Amaral, 1978), but subsequent studies have found no differences in morphology or connectivity between CA4 and CA3 neurons and thus contemporary definitions consider CA4 as part of the CA3 field (Amaral et al., 2007; Maliković et al., 2022). Lorente de Nó separated the regio inferior, populated by large pyramidal cells into two subfields: CA3, located proximal to the DG and receiving mossy fiber input from granule cells and CA2, a small field bearing pyramidal cells with similar morphology to those in CA3 but lacking mossy terminal input from the DG. Also, he differentiated CA1 as populated by smaller pyramidal cells, resembling the regio superior of Cajal. However, subsequent studies on the hippocampus usually did not recognize CA2 as a separate field, and either overlooked it, or merged it with CA3, in a division that ultimately resembled that of Cajal (Ramon y Cajal, 1893; Schlessinger et al., 1978; Boss et al., 1987; Seress, 1988; Ishizuka et al., 1990). There were several reasons for merging both fields, as some authors considered CA2 as a small transitional field (e.g.: (Gaarskjaer, 1986; Woodson et al., 1989). Also, there were practical reasons, as the morphology of the pyramidal cells in CA2 is very similar to those in CA3 (Woodhams et al., 1993; Ishizuka et al., 1995) and it is difficult to separate them. Ultimately, it was reasoned that adding CA2 to field CA3 would not affect significantly the quantification of neurons in CA3, given the small size of CA2 (e.g.: (Boss et al., 1987). This practice of merging CA3 and CA2 has become standard, and we found only 4 studies reporting on CA3, and 2 studies quantifying CA2. In rare cases, reports on CA3 do not specify if they are referring to CA3 or CA3 and CA2 combined, so we classified those estimates to the best of our knowledge based on the figures or descriptions provided by the authors. Regarding CA2, we found 2 quantifications, providing a rough estimate that lacks reliability. Despite this scarcity, it seems relevant to consider CA2 separately, as recent research has revealed its unique molecular profile and connectivity, setting CA2 clearly apart from their neighbors (Middleton and McHugh, 2020). Hopefully, this renewed interest will lead to more stereological descriptions of its neuronal population in the future.

Regarding the rest of the fields, the subicular complex is normally parcellated in three regions: presubiculum, subiculum and parasubiculum, although some studies (e.g.: (Amaral et al., 2024)) further differentiate the postsubiculum, corresponding to the most dorsal part of the presubiculum (Swanson and Cowan, 1977; van Groen and Wyss, 1990). The entorhinal cortex is divided in rodents in a medial domain (medial entorhinal cortex: MEC) and a lateral domain (lateral entorhinal cortex: LEC) (Witter et al., 2017).

### Methodology

The articles reported here were part of a collection on the subject by the authors, expanded by comprehensive online searches using keywords such as “rat AND hippocampus AND stereology” and further search of articles from the labs publishing those studies and from their references, that resulted in an initial collection of 166 studies reporting estimates of the number of neurons in the rat hippocampus. Detailed review of the methodology and the results produced a selection of 87 publications. We selected only studies using design-based stereology, derived from modern stereological definitions, as they are expected to render more precise and accurate estimations than older assumption-based studies whose accuracy will depend heavily on the accuracy of the assumptions (for detailed discussion see (Slomianka, 2021)). Design-based stereology includes Nv*Vref schemes, where the number of neurons is calculated as the product of the density of neurons (Nv) and the volume of the structure under analysis, also called reference volume (Vref). Neuronal density is calculated with the dissector method, either optical or physical, and the volume is obtained with the Cavalieri method (Slomianka, 2021). The Nv*Vref scheme has been mostly replaced in the last decades by the optical fractionator method, that uses the optical dissector to obtain the number of neurons in a known fraction sample of the region of interest, that can then be extrapolated to the whole structure. This method waives the need to calculate Vref, avoiding one source of variance in the estimate.

Additionally, we reviewed the material and methods and selected studies using naïve animals and sham controls from experimental conditions, as the variability did not differ in both populations, but excluded those using special strains (Chen et al., 2006; Kaae et al., 2012) or controls subjected to treatments that could produce neuronal death, like large accumulated doses of BrdU <u>(e.g.: Olesen et al., 2015; Alemu et al., 2019)</u>). Also, we excluded studies performing partial quantification of the hippocampus (e.g.: only septal: (Lemaire et al., 1999; Karimi et al., 2018; Kubová et al., 2018)) and those quantifying neurons in all strata (not restricted to principal cells in the GCL or stratum pyramidale). Regarding stereological methodology, we excluded studies using nucleoli as the counting particle, as is known that hippocampal neurons in the rat frequently exhibit more than one nucleolus (Gaarskjaer, 1978; Bayer et al., 1982; Seress, 1988), except when they acknowledged they counted only one per neuron (Zhao et al., 2016). Also, we excluded studies that suggested the use of partial dissectors; studies that introduced unnecessary shrinkage correction, or Nv*Vref studies that calculated neuronal density and volume in different tissue, as it introduces an additional source of variance. Additionally, we excluded studies reporting higher number of neurons in CA3/CA2 than in CA1, as it contradicts most available descriptions and suggests unconventional segregation of CA fields. Finally, unrealistic low and high values were excluded (e.g.: estimates of 74,000 or 1,300,000 neurons in CA1). For further details see **Suppl. Table 1.**

Estimates of number of neurons correspond to the cellular stratum or principal cell layer of each field, meaning GCL of the dentate gyrus, stratum pyramidale of CA fields and subiculum, and layers II to VI of the entorhinal cortex. In the case of the hilus (the polymorphic layer of the DG), we report all hilar neurons, excitatory and inhibitory, as is typically described in quantitative studies.

For each estimate, we recorded the methodology used, number of animals, strain, sex and age at sacrifice of the subjects. The definitions of life stages follow previously published data from our lab (Arellano et al., 2024). In cases where age was not reported and body weight was provided instead, we estimated age using growth charts for SD (Brower et al., 2015) and Wistar rats (Xie and Zaidi, 2013). Neuron counts are reported per hemisphere, as is standard in the literature. When studies reported the total number of neurons in both hemispheres (e.g.: (Amaral et al., 2024), values were halved. Most reports do not indicate if estimates are uni- or bilateral and in those cases, we have assumed data is per hemisphere. In a few cases, separate numbers for left and right hemispheres are provided (e.g.: (Baldwin et al., 1997; Lister et al., 2006)) and the average of both hemispheres was listed. Some studies reported data on mixed males and females that is included in the tables to illustrate the overall distributions but have been excluded from further analysis given their small number and the potential for sex dimorphism in the number of neurons.

The mean is used to obtain average values of number of neurons in each field, but, when possible, the median is also provided, as an alternative estimate of the average value that is less affected by outliers or skewed data. The variability of the estimates was estimated using the coefficient of variation CV), corresponding to the standard deviation divided by the mean. Outliers were identified using the Tukey method (Tukey, 1977): values below quartile 1 (Q1) – (1.5 × interquartile range (IQR)) or above Q3 + (1.5 × IQR) were excluded from statistical analyses. Normality was verified with the Shapiro-Wilk test, allowing comparisons with the Welch t-test, a variant of the Student’s t-test that requires normal distribution but assumes unequal variances or uneven sample sizes. Statistical significance was set at p<0.05. The minimum number of estimates required for statistical analysis is 5. For completeness, averages obtained from less than 5 estimates are shown in tables, but they are italicized to indicate limited reliability.

## Results

We have analyzed 87 studies documenting a total of 264 stereological estimates of number of neurons in the DG, CA fields, subicular complex and entorhinal cortex of rats, as detailed in **Table 1**. About 80% of the studies used the optical fractionator method and rendered 85% of the estimates, with the remaining 20% of studies using Nv*Vref schemes (see methods) and generated 15% of the estimates. The studies used 4 strains of rats, although 83% of the estimates originate from Wistar (Wi; 46%) and SD (SD; 37%) with marginal contribution of data from Fisher 344 (F344; 11%) and Long Evans (LE: 6%) rats.

**Table 1.** Summary of the sex, strain and age of the animals in the collected studies. Total studies were 87, but the sum of percentages exceeds 100% because some studies analyzed animals at different ages, from two strains or both male and females. Age intervals defined according to (Arellano et al., 2024): adolescence (Adol.) starts at 0.75 months; young adulthood (Y. adult) starts at 2 months; middle age (Mid. age) starts at 6 months, and old age starts at 15 months. SD: Sprague-Dawley; Wi: Wistar; F344: Fisher 344; LE: Long-Evans.

| Sex | % studies | % param | Strains | % studies | % param | Age | % studies | % param |
| --- | --- | --- | --- | --- | --- | --- | --- | --- |
| M | 74 (85) | 209 (79) | Wi | 38 (44) | 122 (46) | Infant | 4 (5) | 6 (2) |
| F | 12 (14) | 45 (17) | SD | 38 (44) | 97 (37) | Adoles. | 14 (16) | 40 (15) |
| M-F | 3 (3) | 10 (4) | F344 | 6 (7) | 28 (11) | Y. adult | 51 (59) | 120 (45) |
|  |  |  | LE | 6 (7) | 17 (6) | Mid.age | 26 (30) | 60 (23) |
|  |  |  |  |  |  | Old age | 12 (14) | 38 (14) |
|  | 89 | 264 |  |  | 264 |  | 107 | 264 |
| Total | (102) | (100) |  | 88 (101) | (100) |  | (123) | (100) |

Regarding sex, almost 80% of the estimates come from males while only 17% are from females and the remaining 4% from mixed males and females. In terms of age, 83% of the estimates come from adult animals: 45% from young adults, 23% from middle-aged and 14% from old animals. Developing animals, including infants and adolescents, accounted for the remaining 17% of the estimates (**Table 1**).

The summary of the findings regarding updated neuronal populations is shown in Fig. 1. The next sections will analyze in detail the findings in each field of the hippocampal formation.

**Figure 1.**
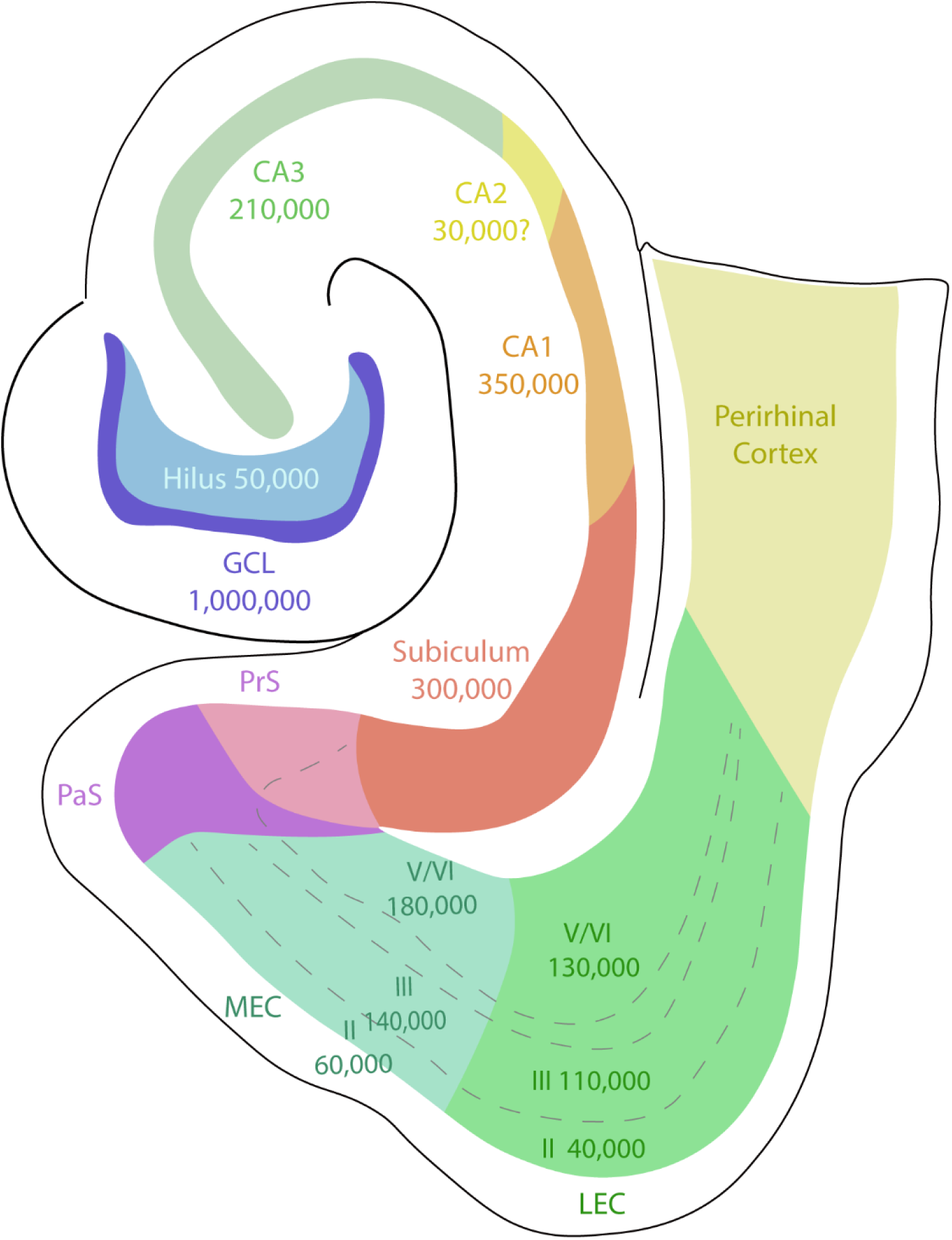
Schematic drawing of the hippocampal formation of the rat (horizontal view) showing the average neuronal populations obtained in the present analysis. Data correspond to adult males except in the EC, where females (adult and adolescent) are averaged (see text). Presubiculum (PrS) and parasubiculum (PaS) are not annotated given the potentially inaccurate averages (see text). Modified from (Cappaert et al., 2015)

### Granular cell layer

We found 42 reports providing 53 estimates of the number of neurons in the granular cell layer (GCL) (**Suppl**. **Table 2)**. About 80% of the estimates come from adult animals, with the rest providing information on developing animals up to two months of age. Most of those adult estimates are from males (87%) and only one estimate is from mixed males and females. The data involved 4 strains, although most estimates originate from Wistar or SD rats (Figure 2). The average number of neurons segregated by age, strain and sex is summarized in **Table 2**. Tukey fences (Tukey, 1977) were 0.44 and 1.6, revealing 1 outlier (1.86 million) that was excluded from further analysis (**Suppl. Table 2**). Male average without outliers was 1.05 million with a similar median of 1.09, suggesting quite symmetric distribution of the data. Female data was scarce, with only 5 datapoints that produced a similar average of 1.01 million GCs, with a median of 1.02. Total male/female average was 1.03 million GCs.

**Figure 2.**
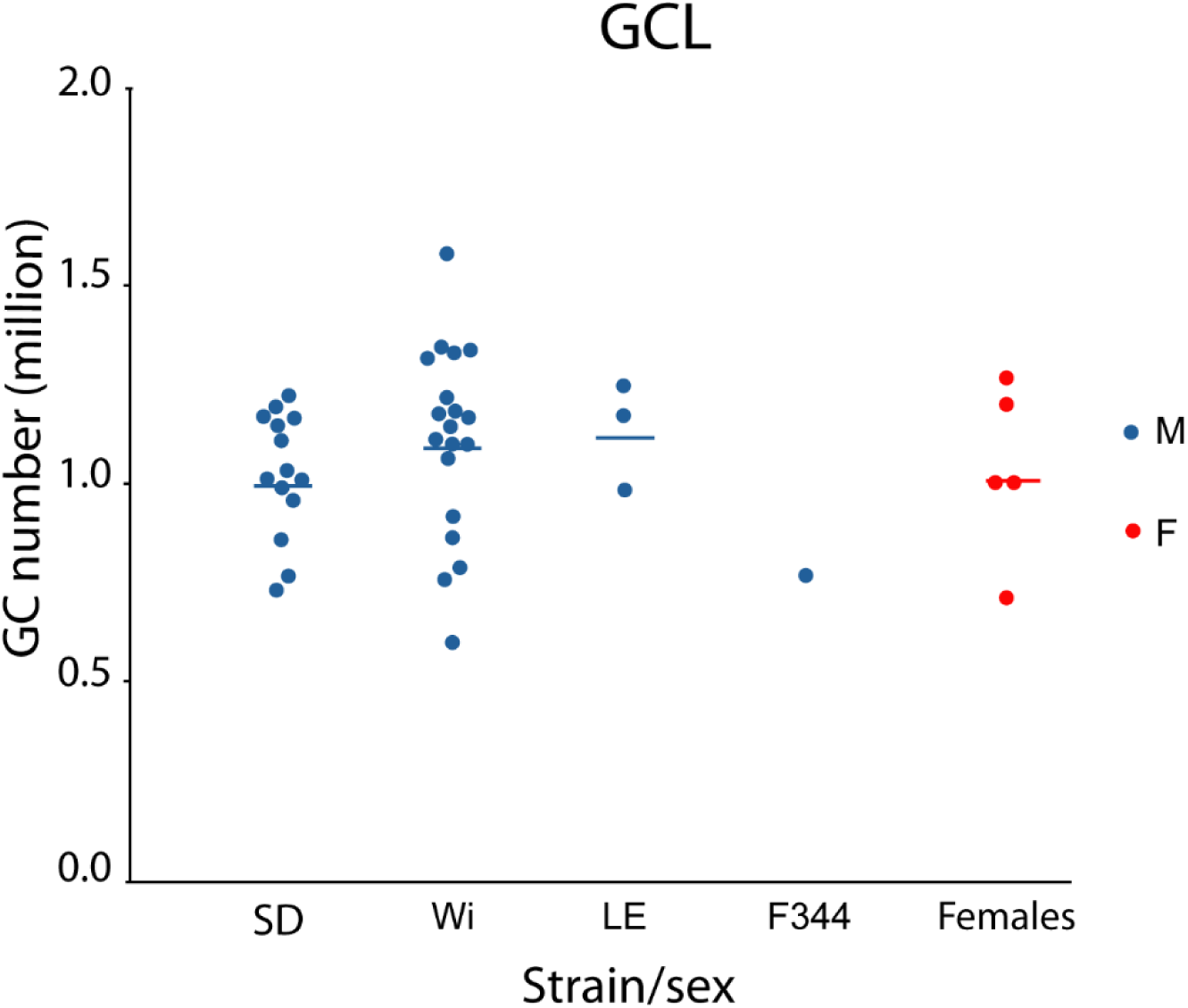
Cloud representation of adult males and females by strain. Averages are represented by horizontal lines. Averages of SD and Wistar males and overall females orbit around ∼1 million GC. Outliers are not represented. GC: granule cells; SD: Sprague-Dawley; Wi: Wistar; LE: Long Evans; F344: Fisher 344; M: males; F: females.

**Table 2:** Comparison of GC numbers in SD and Wistar males and in total males and females for both adolescent and adult animals. Average for adolescent SD males and adolescent females might not be reliable (n<5) and are highlighted in italics. Total males and females include all strains; SD: Sprague-Dawley; CV: coefficient of variation; n: number of estimates.

| Age | Males |  |  | Females |
| --- | --- | --- | --- | --- |
|  | SD (CV, n) | Wistar (CV, n) | Total (CV, n) | Total (CV, n) |
| Adolescent | *0.95 (0.23, 2)* | 0.98 (0.19, 5) | 0.97 (0.18, 7) | *1.04 (0.22, 2)* |
| Adult | 0.99 (0.17, 14) | 1.09 (0.22, 19) | 1.05 (0.20, 37) | 1.01 (0.22, 5) |

### Age differences

Graphic representation of the data across age **(Figure 3)** shows large dispersion of the data at every age. There was a mild increase in neuron number early postnatally that stabilizes after age 2-3 months, reflecting granule cell neurogenesis that occurs mostly after birth, with a peak around the end of the first postnatal week (Schlessinger et al., 1975; Bayer, 1980a) and is followed by a sharp decrease during adolescence to continue at low levels during adulthood (Arellano and Rakic, 2024). Adolescent (0.75-2 months of age) males (n=7) showed 0.97 million GCs, quite close (8% difference) to the number in adults (Mann-Whitney U test, p=0.41). There were only 2 adolescent datapoints for females (average 1.04 million) that precluded meaningful comparisons.

**Figure 3.**
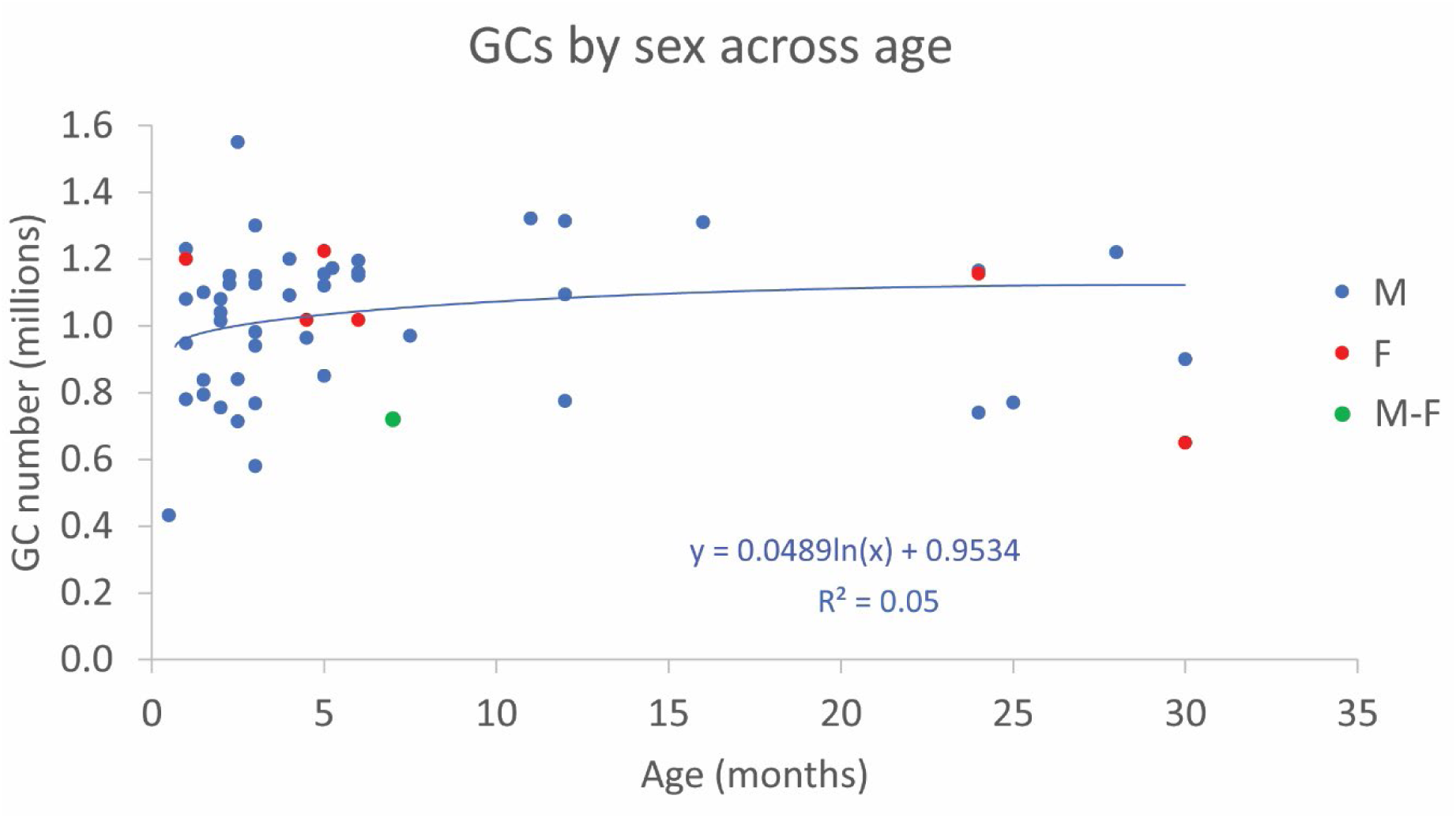
Distribution of datapoints across age (excluding outliers). Note the dispersion of the data across ages that compromises regression (R2=0.05). No clear differences can be observed between the distribution of female (red) and male (blue) data. There was only one datapoint for mixed males and females (green).

The presence of adult neurogenesis in the granular cell layer invites two potential scenarios: new neurons replace existing ones, resulting in a stable population, or new neurons are added to the existing population, producing an increase in the number of granule cells with age. A third possibility is a mix of both scenarios, resulting in a modest increase in the GC population over time. Several studies have provided data regarding the number of GCs at different ages. Studies focused on developing animals typically show an increase in GCs, as expected (Duffell et al., 2000; Heine et al., 2004; Smith et al., 2008; Elibol-Can et al., 2014). However, in adults, only two studies one ABS study (Bayer et al., 1982; Rasmussen et al., 1996) reported clear increase in the number of GCs across age, while many other studies have not found significant increases (Andrade et al., 1995, 1996; Brandão et al., 1995; Rapp and Gallagher, 1996; Merrill et al., 2003; Heine et al., 2004), suggesting that either few new neurons are added to the adult population, or new neurons might replace existing ones. Modelling of adult neurogenesis in rats indicates that only about 120,000 new neurons are added to the GC during adulthood, between age 2 months and 21 months (Arellano and Rakic, 2024), a small increase that cannot be detected with stereological quantifications, as they describe the overall population dynamic with broad strokes, as illustrated in **Figure 3**.

The presence of adult neurogenesis in the granular cell layer invites two potential scenarios: new neurons replace existing ones, resulting in a stable population, or new neurons are added to the existing population, producing an increase in the number of granule cells with age. However, modelling of adult neurogenesis in rats indicates that only about 120,000 new neurons are added to the GC during adulthood, between age 2 months and 21 months (Arellano and Rakic, 2024), a small difference that likely will not be detected with stereological estimates, as they do not provide enough resolution. In this regard, developmental studies typically show an increase in GCs, as expected given the mostly postnatal development of the GCL (Duffell et al., 2000; Heine et al., 2004; Smith et al., 2008; Elibol-Can et al., 2014). In adults, however, although (Bayer et al., 1982) using assumption-based stereology and (Rasmussen et al., 1996) using the optical fractionator reported clear increase in the number of GCs across age, several others did not find significant increases (Andrade et al., 1995, 1996; Brandão et al., 1995; Rapp and Gallagher, 1996; Merrill et al., 2003; Heine et al., 2004), suggesting that the population of GCs is essentially stable across adulthood, or might experiment small variations that might run undetected by stereological studies. In this regard, the lack of clear regression (R2<0.1) of our dataset precludes any reliable modelling of those potential increases in the GC population across age.

### Strain differences

Previous studies using ABS have shown strain differences in the number of GCs. (Boss et al., 1985) found that adolescent SD females had about 45% more GCs than SD. In the same line, (Seress, 1988) reported that mixed male and female adult SD rats showed 35% more GCs than Long Evans rats. We compared only adult Wistar and SD males, as there were only scarce data on other strains, from females or from adolescent SD males (**Table 2)**. Our results showed 1.09 million GCs in adult Wistar males and 0.99 million in adult SD males (**Table 2**), a 9% difference that was not significant (two tailed Welsch test, p=0.16) but shows an opposite trend to that reported by (Boss et al., 1985).

### Sex differences

Several studies have reported sex differences in GC numbers (Madeira et al., 1988, 1991; Nuñez and McCarthy, 2003; Nuñez et al., 2003a, 2003b; Schmitz et al., 2005; Smith et al., 2008; West et al., 2021). All but one of those reports (West et al., 2021), showed that females exhibit 9-28% less GCs than males both during early development in the first postnatal month (Madeira et al., 1988; Nuñez and McCarthy, 2003; Nuñez et al., 2003b, 2003a; Smith et al., 2008) or in adult animals (Madeira et al., 1991; Schmitz et al., 2005; Smith et al., 2008; West et al., 2021), suggesting the presence of sex differences in the number of GCs in the dentate gyrus. Comparison between males and females in our dataset has limited reliability due to the scarcity of adult female datapoints (n=5) but shows no clear differences (3% less GCs in females; one tailed Welsch test p=0.38; **Table 2**).

Data collected here show an overall average of 1.03 million GCs, matching well the traditional 1 million value used in many descriptions and models of the rat hippocampus. We could not reveal clear differences between males and females, although the female dataset is very limited. We found a 9% difference between Wistar and SD strains (males) that was however not significant, as the samples exhibit large variability.

### Hilus

The hilus corresponds to the polymorphic layer of the DG and is populated by excitatory mossy cells and GABAergic interneurons. We selected 38 studies that produced 42 estimates of the total number of neurons in the hilus, mostly from adult males, with scarce data from adolescents (n=4 males, 1 female) or adult females (n=3), that precludes meaningful comparisons by sex of between adolescent and adults **(Figure 4**, **table 3)**. Outlier analysis by Tukey fences (29,721 and 67,457) revealed two outliers (68,000 and 115,000) that were excluded from the analysis (**Suppl. Table 3**). Focusing on adult males, the hilus showed an overall average of 48,000 neurons and a close median of 49,000 neurons, suggesting symmetrical distribution of data and adequate outlier detection. Analysis by strains did not reveal differences between Wistar and SD, both exhibiting ∼48,000 neurons, very similar to the average of 49,000 in Fisher 344 males, although this is based in only 3 estimates and might not be reliable. (Xu et al., 2004) compared Wistar and Long Evans adult males and reported similar values of 43,000 and 45,000 neurons respectively, supporting the absence of strain differences in the hilus (**Figure 4**, **Table 3**).

**Figure 4.**
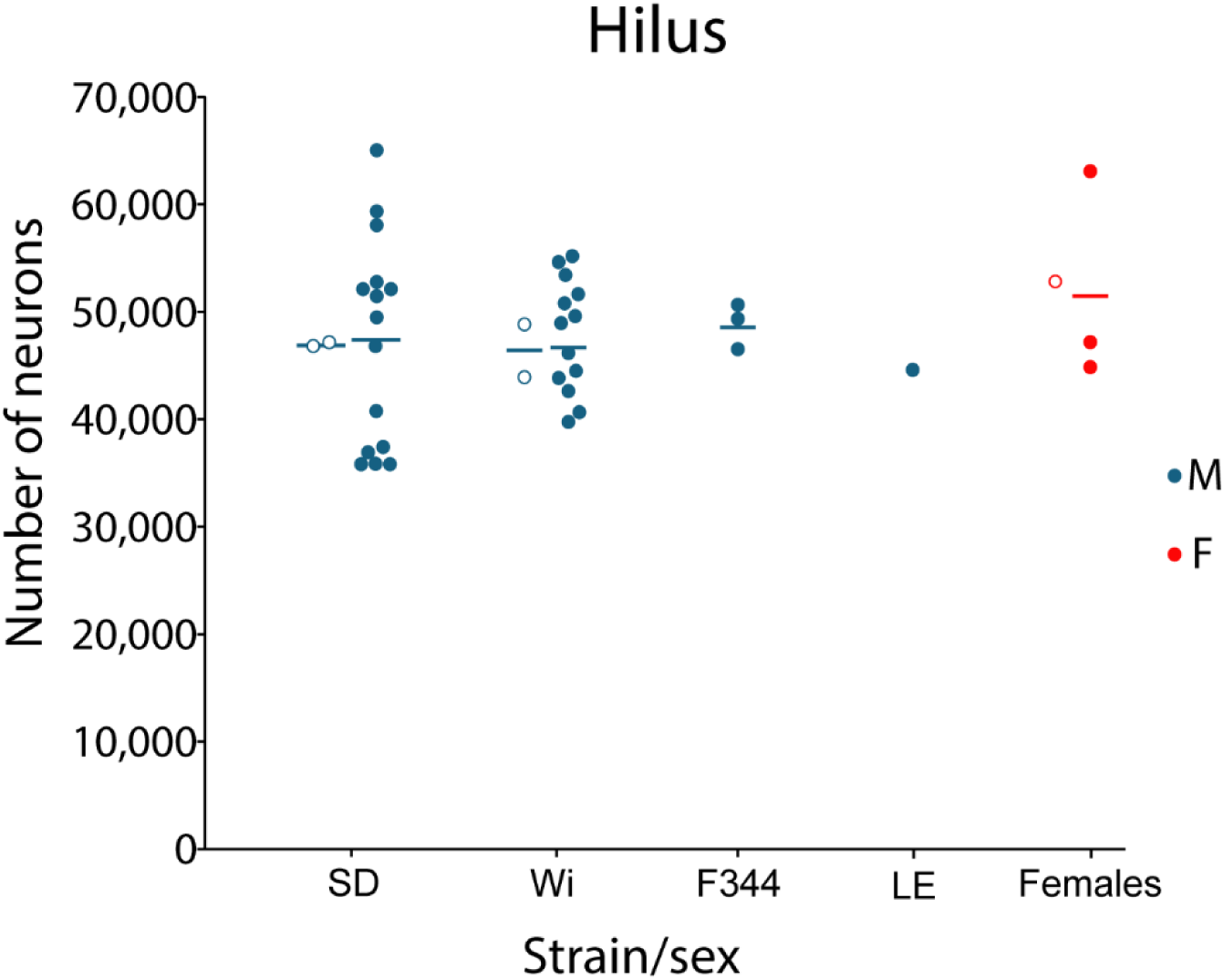
Number of neurons in the hilus by strain and sex. Circles represent adolescent values and filled dots represent adults. Averages are represented by horizontal lines. For simplicity, female data is pooled from 3 strains.

**Table 3:** Average number of hilar neurons in adolescent and adult males and females and Wistar and SD males. Averages for adolescent animals and adult females are not reliable (n<5) and are highlighted in italics. Estimates were quite homogenous across age and sex. Total adult males and females include all strains. SD: Sprague-Dawley; CV: coefficient of variation; n: number of estimates.

|  | Males |  |  | Females |
| --- | --- | --- | --- | --- |
|  | SD (CV, n) | Wistar (CV, n) | Total (CV, n) | Total (CV, n) |
| Adolescent | *47,150 (0.004; 2)* | *46,560 (0.08; 2)* | *46,855 (0.04; 4)* | *53,000 (1)* |
| Adult | 47,626 (0.21; 15) | 48,097 (0.11; 13) | 47,856 (0.16; 32) | 51,733 (0.19, 3) |

Three studies provided data across adult ages that showed no clear differences between young and old individuals (Rasmussen et al., 1996; Merrill et al., 2003; Hattiangady et al., 2005). Only (Rasmussen et al., 1996) described a 20% decline in hilar neurons between age 2.5 and 24 months, but the other two studies showed stable or slightly increasing (5%) populations, and therefore it seems the population of hilar neurons is stable across the lifespan. The data in our collections shows large variability across ages without clear regression (**Suppl. Fig 1**).

Overall, both the average and the median suggest the hilus has about 50,000 neurons in males, with similar estimates across strains. There are not enough datapoints in adolescents or females to reach reliable averages, and therefore no comparisons can be established by sex or age.

### CA3-CA2

As mentioned above, most studies on CA3 include also CA2 (for details see material and methods), resulting in a combined field CA3-CA2. We found 28 studies that produced 34 estimates of the number of neurons in the stratum pyramidale of CA3/CA2, mostly from adult males (n=26) and only 2 estimates from females (**Suppl. Table 5, Table 4 and Figure 5**). Tukey fences for adult animals (100,068 and 373,888) detected 2 outliers that were excluded from further analysis.

**Figure 5.**
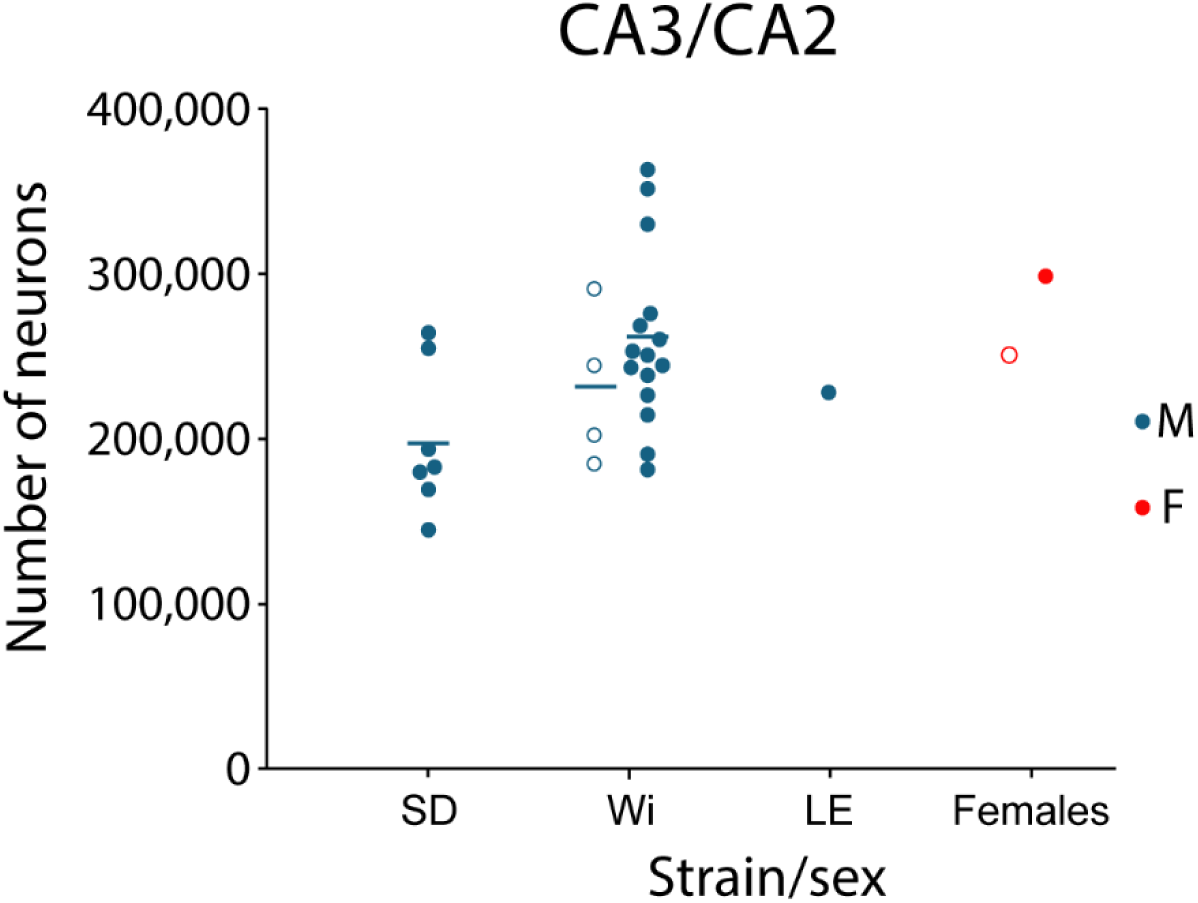
Cloud graph showing the estimates of number of neurons in CA3/CA2 per strain and sex. Circles represent adolescent values and filled dots represent adults. Averages are represented by horizontal lines. Note that female information is scarce and only available for SD and Wi strains.

**Table 4:** Average of CA3-CA2 neurons per strain in adolescent and adult males. Adolescent data and adult female data are not reliable given the low number of estimates and are highlighted in italics. Total adult males include all strains. SD: Sprague-Dawley; NA: no data available; CV: coefficient of variation; n: number of estimates.

| Age | Males |  |  | Females |
| --- | --- | --- | --- | --- |
|  | SD (CV, n) | Wistar (CV, n) | Total (CV, n) | Total (CV, n) |
| Adol. | NA | 229,992 (0.21, 4) | 229,992 (0.21, 4) | 250,000 (1) |
| Adult | 197,060 (0.22, 7) | 257,389 (0.21, 15) | 237,219 (0.23, 24) | 298,000 (1) |

The data shows, as in the GCL, a considerable degree of variability across age (**Suppl. Fig. 2**) that precludes regression (R2=0.1). There are only 4 estimates for adolescent males, showing an average of 230,000 neurons, similar (3% difference) to the average of 242,000 neurons for adult males (**Table 4**).

Contrary to granule cells that exhibit protracted postnatal neurogenesis, neurons in the rest of hippocampal fields are generated largely pre- and perinatally, reaching a peak around the end of the first postnatal week and then pruning to reach adult numbers around P15 (Bayer, 1980b, 1980a; Bandeira et al., 2009). As differentiation proceeds during those postnatal weeks, it seems reasonable to assume that by the onset of adolescence, at around 3 weeks of age (Arellano et al., 2024), the number of cells identifiable as neurons would match adult levels. However, some previous studies have shown that adolescent animals exhibit consistently less neurons (average 89%; range 83-92%) than adults, with similar percentages for males and females (Madeira et al., 1992) in CA fields (Madeira et al., 1992; Wakuda et al., 2008; Partadiredja and Bedi, 2010; Elibol-Can et al., 2014), with the only exception of (De Araujo Furtado et al., 2024), that reported similar number of neurons (within 5%) between adolescent and adults in CA1-CA2. In our dataset, we have only scarce data from adolescents that preclude establishing reliable comparisons, but the average of the few datapoints available for adolescents is similar (3% lower) to the adult average, in line with the results in the hilus.

When comparisons are established by strains (**Figure 5**), we focused on SD and Wistar, as there are only 2 datapoints for adult Long Evans males (average 227,000 neurons). Adult SD males showed 23% less neurons than Wistar, a remarkable difference that reached statistical significance (p=0.014; two-tailed Welsch test). At face value, the large difference between adult SD and Wistar males seems reliable, as is based in 7 and 15 estimates respectively. However, the dataset also shows that adult Wistar males contain two high values (350,000, 360,000) coming from the (Rasmussen et al., 1996) study that also produced high estimates in CA1 (that were detected as outliers), while SD males contain data from studies that also report low estimates in other fields, like (Chen et al., 2008; Keleş et al., 2019). Thus, the large difference detected might be inflated by those studies’ influence. Indeed, if those low and high estimates mentioned above are removed, we obtain averages of 213,000 for SD males (n= 5) and 242,000 for Wistar males (n=13), with a more moderate difference of 12%. However, there are no objective reasons in the methodology or results of those studies to be excluded from the analysis, and so we reported them. Indeed, the main goal of this analysis is to obtain more balanced averages that might compensate for high or low values produced in individual studies. This goal might not be achieved in every case, particularly when the number of datapoints is relatively low, as in this case (n=7). In that regard, the average of all adult males might buffer those potential low or high averages and produce a more accurate estimate for the overall population.

### Number of neurons in CA3 alone

The neuronal population of CA3 alone in males can be estimated by subtracting the population of CA2 from the total of CA3/CA2. As described in the CA2 section, it might represent 12.9% of CA3-CA2, or about 30,000 neurons (12.9% of 237,000) in adult males, meaning that CA3 would represent ∼210,000 neurons (∼87% of 237,000; **Suppl. Table 5, 6)**. That value is consistent with the average of 215,000 obtained from studies quantifying CA3 alone (**Suppl. Table 4**), and thus it seems reasonable to assume that CA3 alone might have ∼210,000 neurons.

### CA2

As indicated above, almost all studies describe pooled data for fields CA2 and CA3. And although we found 10 studies that quantified the number of neurons in the stratum pyramidale of CA3 and CA2 separately (**Suppl. Table 6**), all of them were excluded from our analysis for varied reasons. (Cassell, 1980) used assumption-based stereology to describe ∼12,400 neurons in CA2, an unlikely low estimate that he reported represented about 8% of CA2/CA3 as described in (Amaral et al., 1990). This study was excluded because of its unreliable methodology (see methods). Another study used the optical fractionator and reported ∼35,000 neurons in CA2 (Wang and Gondré-Lewis, 2013) but was excluded because it used nucleoli as the counting particle (see methods). Three other studies (Golub et al., 2015; Reddy et al., 2024; Singh et al., 2024), reported 71,000-78,000 neurons for total neurons in all CA2 strata. Although those figures are useful as a reference, they cannot be compared with data restricted to principal neurons in the stratum pyramidale and were excluded from the analysis. Another study by (Köylü et al., 2021) using an Nv*Vref scheme with physical dissectors provided a similar estimate of 72,000 neurons in adult males, representing ∼27% of CA3/CA2 neurons. Both the estimate and the percentage seem very high and based on the coincidence with the studies describing all CA2 neurons it was also excluded from the analysis. Finally, four other studies, all from the same laboratory, were excluded as they provided exceedingly large numbers from 133,00 to 650,000 neurons (Yurt et al., 2018; Deniz and Kaplan, 2022; Elamin et al., 2022; Hamadi et al., 2022) (**Suppl. Table 6**).

With all those caveats in mind, it seems that we do not have reliable figures to estimate the number of pyramidal cells in CA2. One possibility is that beyond the large differences in absolute number of neurons, those studies report similar proportion of CA2 neurons compared to CA3/CA2 or CA1, that would allow obtaining an estimate of CA2 out of our estimates of those populations. Using data from fields CA3 and CA1 in those studies, it can be calculated that CA2 represents 8-46% of CA3/CA2 and 6-71% of CA1, suggesting that beyond quantification differences, they might also differ markedly in the segregation of CA fields.

As an alternative rough estimate, we measured the relative volume of CA1, CA2 and CA3 in the mouse atlas from the Allen Institute, assuming their volumetric ratios might be similar to those of the rat (**Table 5**). The results showed that CA2 represents about 10% of the volume of CA3 and about 8% of CA1 (**Suppl. table 1**). To compensate for potential differences in neuronal density between those fields, we incorporated actual densities reported by (Wang and Gondré-Lewis, 2013) indicating that neuronal density in CA2 is 1.5 times that of CA3 and 1.2 times that of CA1. Using those ratios, the neuronal population of CA2 would be ∼15% of the population of CA3 or 32,300 and 9.5% of the population of CA1 or 33,100 (**Table 5**). Those estimates agree well with each other, but it can be argued that the CA3 estimate is not reliable, as it is based only on 4 datapoints. However, estimating the proportion of CA2 neurons over the total in CA3/CA2 reveals that it represents 12.9% or 30,600 neurons (12.9% of 237,000), that once again agrees well with the other estimates, and suggests that the number of pyramidal cells in CA2 might be around 30,000 neurons, representing about 1/8 of the CA3/CA2 population of ∼240,000 neurons. Following that logic, CA3 would have ∼210,000 neurons, in good agreement with the estimate of 215,000 neurons for CA3 alone, an estimate that might be more accurate than expected considering it is based on only 4 datapoints. Thus, until more precise data about the neuronal population of CA2 is available, it seems reasonable to assume that CA2 might have ∼30,000 neurons, or about 13% of the combined population of CA3-CA2.

**Table 5.**
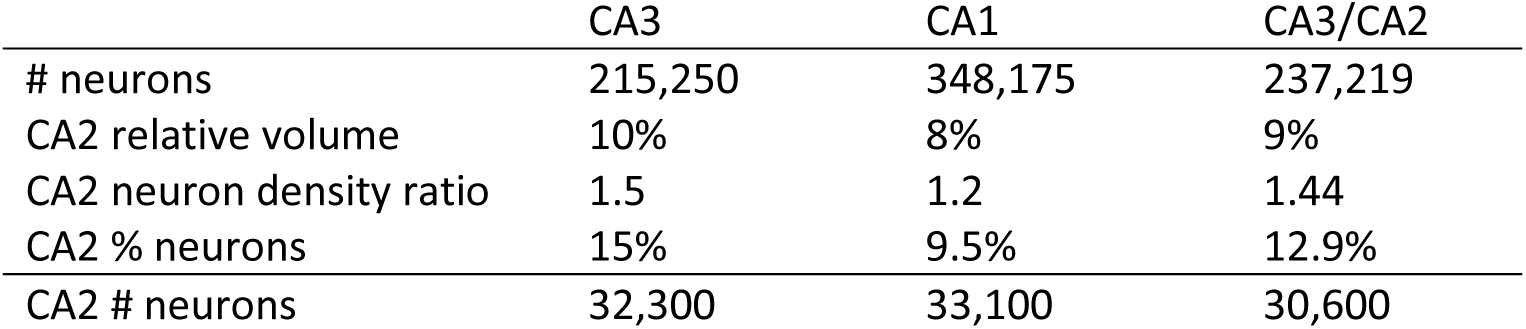
Estimates of the number of neurons in CA2. Estimates are based on the ratio of volume of the stratum pyramidale and neuronal density in CA2 compared to CA3 and CA1. Volume ratios were estimated from the Allen mouse atlas and density ratios from (Wang and Gondré-Lewis, 2013).

|  | CA3 | CA1 | CA3/CA2 |
| --- | --- | --- | --- |
| # neurons | 215,250 | 348,175 | 237,219 |
| CA2 relative volume | 10% | 8% | 9% |
| CA2 neuron density ratio | 1.5 | 1.2 | 1.44 |
| CA2 % neurons | 15% | 9.5% | 12.9% |
| CA2 # neurons | 32,300 | 33,100 | 30,600 |

### CA1

We obtained 42 studies documenting 48 estimates of number of neurons in the stratum pyramidale of CA1, mostly from males (n=45), with only 1 estimate from adolescent females and 2 from mixed male and females, thus precluding comparisons across sexes (**suppl table 7 and Figure 6**).

**Figure 6.**
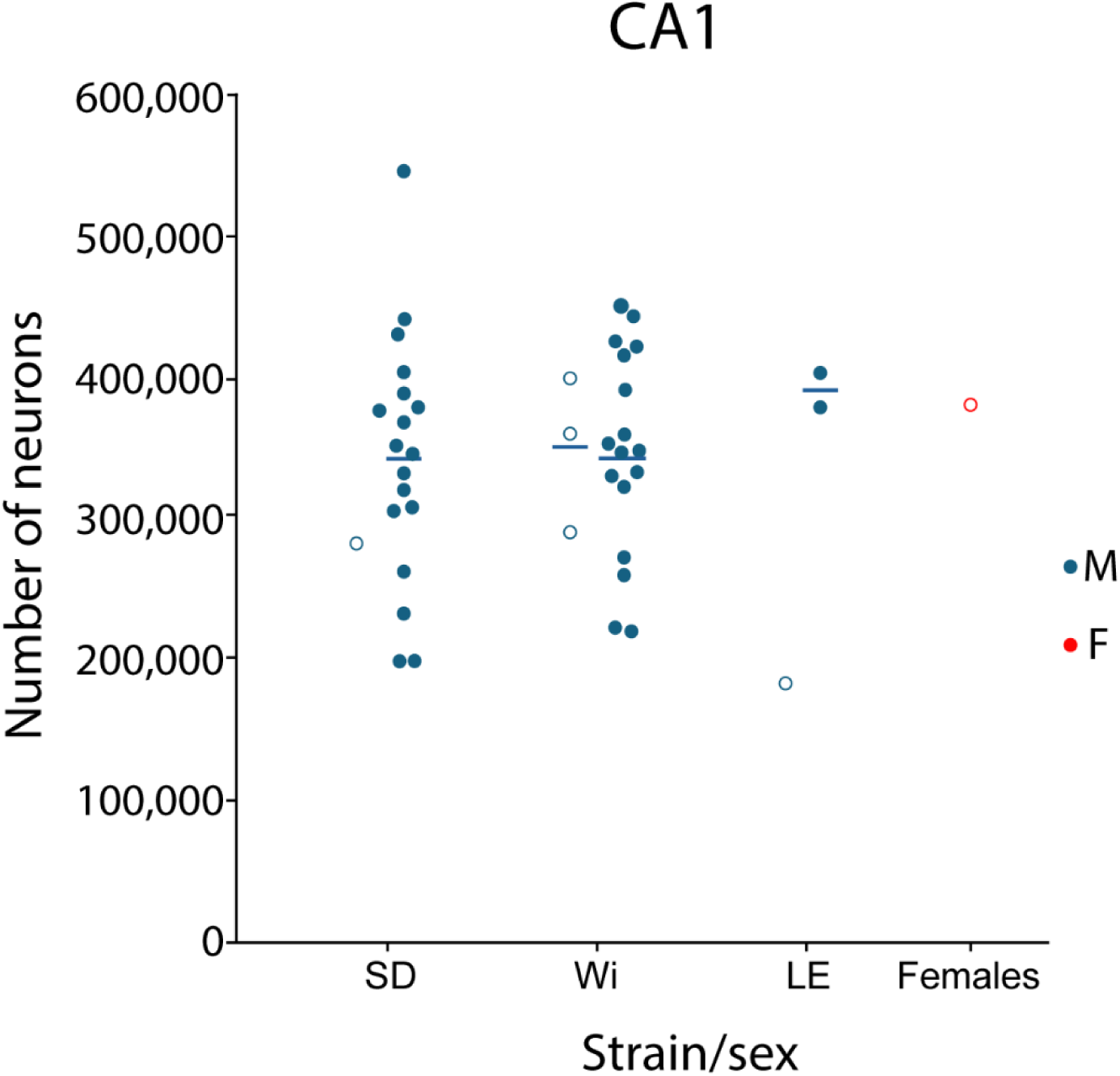
Number of neurons in CA1 by strain and sex. Circles represent adolescent values and filled dots represent adults. Averages are represented by horizontal lines. There is a high value (545,375 in SD) that were however not detected by Tukey fences (169,299 and 556,997). Only one adolescent female datapoint was found. LE: Long-Evans; SD: Sprague-Dawley; Wi: Wistar; M: male; F: female.

Tukey fences (169,299 and 556,997) revealed two outliers (570,000 and 670,000 from (Rasmussen et al., 1996) that were excluded from the analysis (**Suppl. Table 7**). Adult males had on average 348,000 neurons in CA1, with a close median of 352,000 and also similar counts for Wistar (348,000) and SD (344,000). There were scarce datapoints for male adolescents (n=4) with an average of 333,000 neurons, that its quite close to the adult value (within 5%) (**Table 6**). Analysis of the estimates across age showed, as in the other fields, no detectable regression (**Suppl. Figure 3**)

**Table 6:** Average of CA1 neurons per strain in adolescent and adult males and females. The average of adolescent females and SD and Wistar males is not reliable, as is based on scarce (n=1-3) data and is highlighted in italics. Total adult males include all strains. SD: Sprague-Dawley; CV: coefficient of variation; ND: no data available; n: number of estimates.

| Age | Males |  |  | Females |
| --- | --- | --- | --- | --- |
|  | SD (CV, n) | Wistar (CV, n) | Total (CV, n) | Total (CV, n) |
| Adol. | 282,874 (1) | 349,840 (0.16; 3) | 333,098 (0.17; 4) | 382,000 (1) |
| Adult | 343,549 (0.26; 18) | 348,094 (0.25; 18) | 348,175 (0.23; 37) | NA |

Overall, if we consider all strains, it can be concluded that CA1 might exhibit about 350,000 neurons in males, while there is no reliable data for females or adolescents. (De Araujo Furtado et al., 2024) found no differences between males and females in CA1-CA2 (see below) suggesting there might be no sex differences in this field.

### CA1/CA2

Although rare, we found 6 studies reporting 12 estimates of CA2 and CA1 together (**Suppl. table 8**). (De Araujo Furtado et al., 2024) studied both males and females and described not differences in the estimates, so they pooled the data together. The average of all adult estimates (including pooled males and females) is 308,000 (CV= 0.09, n=8) while the average of adult males was 316,000 (CV=0.13, n=4). If we assume that CA2 has ∼30,000 neurons, then the estimated CA1 average would be ∼278,000 and ∼286,000, that are quite lower than the average of 350,000 obtained from CA1 data.

### Subicular complex

We collected 14 studies reporting 22 estimates on the subicular complex fields, mostly on the subiculum (12 estimates; **Suppl. Table 9)**, while 3 studies reported on the presubiculum, parasubiculum and postsubiculum (**Suppl. table 10**).

### Subiculum

We found 12 estimates, most of them (n=10) in males, orbiting around 300,000 neurons, with an average of 279,000 neurons and a median of 296,000 neurons (**Suppl table 9 and Figure 7**). The lower average is due to two discordant estimates below 200,000 that were however not detected as outliers (Tukey fences: 181,992-390,408). The two remaining estimates from females also were very close to 300,000, strongly suggesting that in this case, the median might be a better estimator of the number of neurons than the average. Regarding strains, SD males have scarce data (n=4) that include the two low datapoints bellow 200,000, lowering the average to 242,000 neurons, 22% less than the Wistar average of 308,000 (**Table 7**). Thus, neither the SD average nor the difference between strains can be considered reliable.

**Figure 7.**
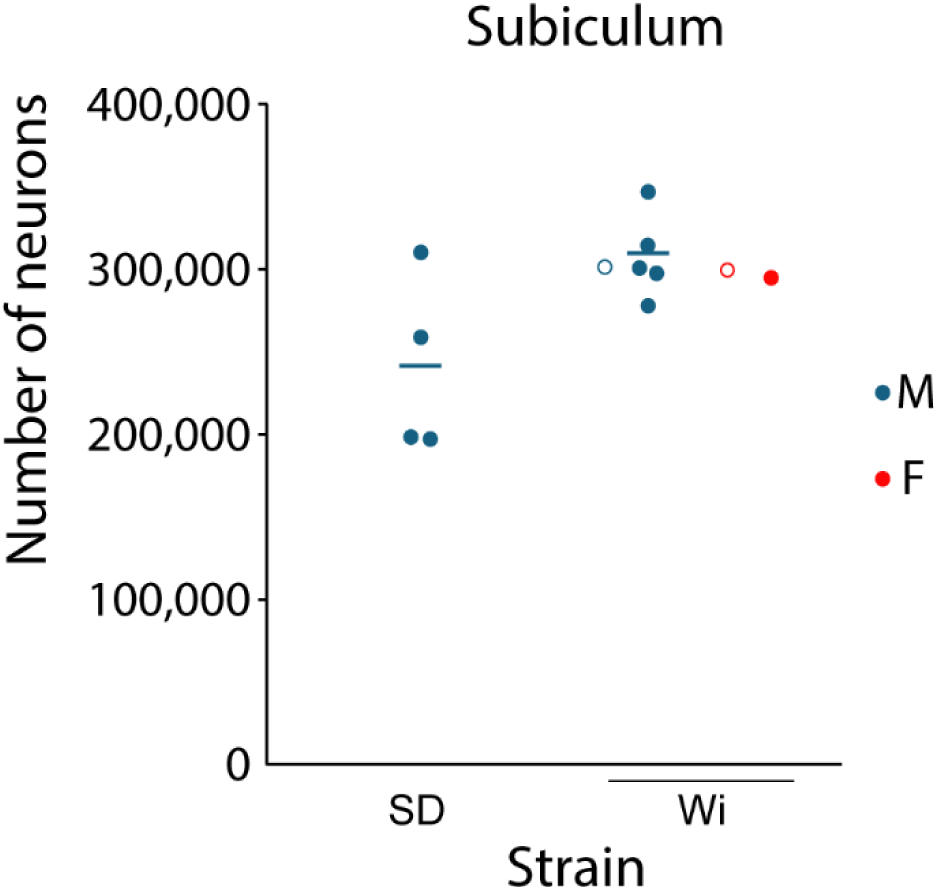
Neuron number in the subiculum per strain and sex. Circles represent adolescent values and filled dots represent adults. Averages are represented by horizontal lines.

**Table 7:** Average of subicular neurons per strain adult males and females. Data per strain and for females are shown for completeness but are not reliable (n=1-4) and are highlighted in italics. Total adult males include all strains. SD: Sprague-Dawley; CV: coefficient of variation; ND: no data available; n: number of estimates.

| Age | Males |  |  | Females |
| --- | --- | --- | --- | --- |
|  | SD (CV, n) | Wistar (CV, n) | Total (CV, n) | Total (CV, n) |
| Adolescent | NA | 303,000 (1) | 303,000 (1) | 300,000 (1) |
| Adult | 242,133 (0.23; 4) | 308,466 (0.09; 5) | 278,985 (0.19; 9) | 296,000 (1) |

The limited data does not allow for reliable comparisons across strain, sex or age, although sex dimorphism, with females exhibiting less neurons than males, has been reported (Andrade et al., 2000). In conclusion, the average for the subiculum is about 280,000 neurons in adult males, while the median is 296,000. Considering the consistency of estimates across sex and age, we think the median might be a better descriptor and can be considered that the subiculum has ∼300,000 neurons, an estimate that is withing the 95% confidence interval for the average.

### Presubiculum, Parasubiculum and Postsubiculum

We found 3 studies reporting the number of neurons in the cellular strata of the pre-, para-and postsubiculum (**Suppl. Table 10**), but their data are not comparable as (Mulders et al., 1997) considered the pre- and parasubiculum, while (Cardoso et al., 2011; Amaral et al., 2024) also differentiated the postsubiculum, a region corresponding to the most dorsal part of the presubiculum (Swanson and Cowan, 1977; van Groen and Wyss, 1990). In addition, (Cardoso et al., 2011) did not differentiate between upper and deep layers in the presubiculum and parasubiculum, while the other studies did. As a result, the results of the 3 studies are listed separately, and no averages are calculated (**Suppl. Table 10)**.

### Entorhinal cortex

The entorhinal cortex (EC) is the main interface of communication between the neocortex and the hippocampus. In murine rodents, the EC is divided into two areas: the medial entorhinal cortex (MEC) and the lateral entorhinal cortex (LEC). In addition, the entorhinal cortex presents 6 layers, although its main feature is that layer IV is basically acellular, populated only with sparse neurons from layer III and V, and is called lamina dissecans, as it separates upper layers II and III from lower layers V and VI (Witter et al., 2017). We analyzed 15 studies that overall contained 43 estimates: some studies segregated the EC by layers (layer II, III and V/VI), some by areas (MEC and LEC), and some presented partial or complete analyses by areas and layers.

### Neuronal number per layer

A modest number of estimates (n=3-5) were available for the number of neurons in layers II, III and combined V/VI of both males and females (**Suppl. Tables 11-13**). The results show females consistently having 11-14% less neurons than males, but neuronal distribution across layers were virtually identical in both sexes, with upper layers taking 53% and deep layers the remaining 47% of the neurons (**Table 8 and Figure 8A**).

**Figure 8.**
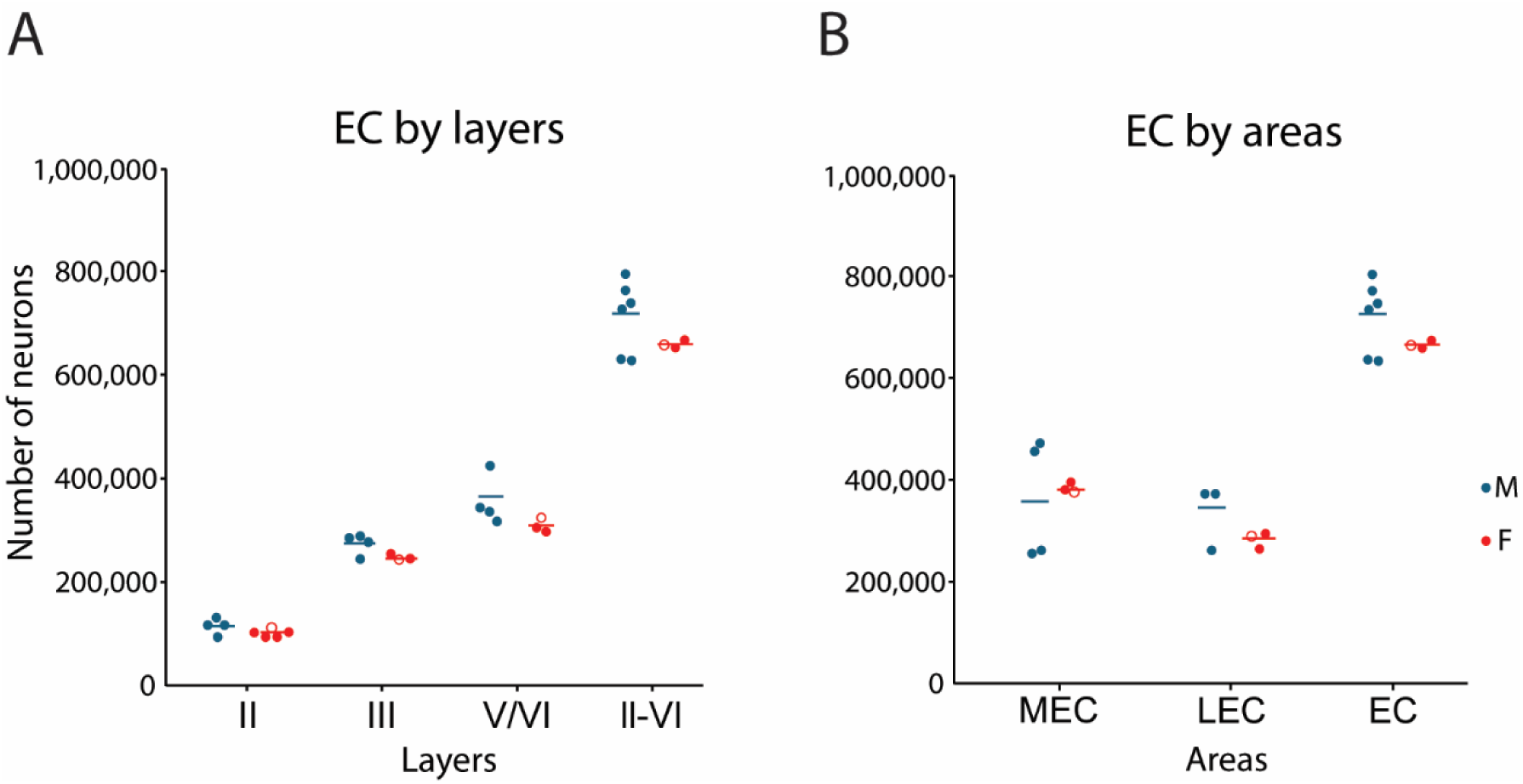
Number of neurons in EC by layers (A) and areas (B). The few female estimates are very homogeneous, while male values are more disperse and tend to show higher averages. Dots represent adults and circles adolescents (n=1). Averages are represented by horizontal lines. MEC: medial entorhinal cortex; LEC: lateral entorhinal cortex; EC: entorhinal cortex.

**Table 8.**
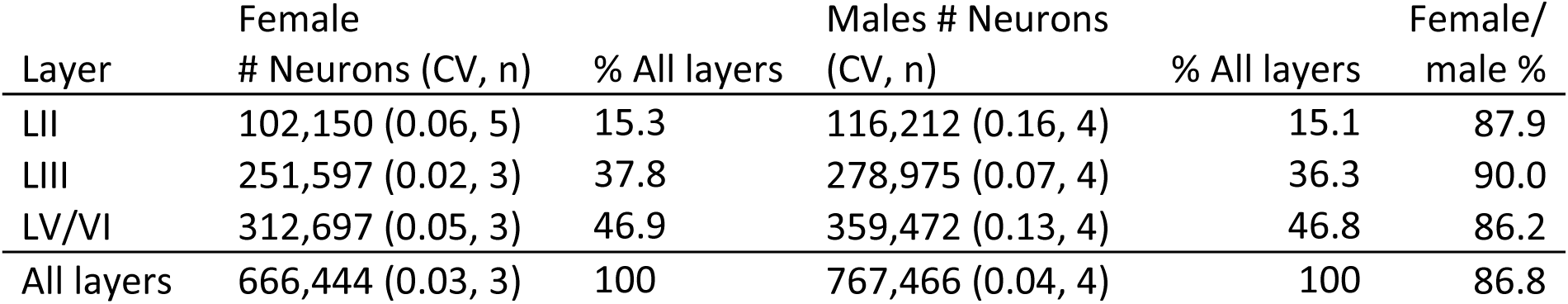
Number of neurons in the EC segregated per layers and sex. The distribution of neurons across layers is virtually identical for both sexes, although females show consistently lower populations. Note: total number of neurons in Layers/areas/EC might vary across tables due to the use of different sets of data to obtain total averages.

| Layer | Female<br># Neurons (CV, n) | % All layers | Males # Neurons<br>(CV, n) | % All layers | Female/<br>male % |
| --- | --- | --- | --- | --- | --- |
| LII | 102,150 (0.06, 5) | 15.3 | 116,212 (0.16, 4) | 15.1 | 87.9 |
| LIII | 251,597 (0.02, 3) | 37.8 | 278,975 (0.07, 4) | 36.3 | 90.0 |
| LV/VI | 312,697 (0.05, 3) | 46.9 | 359,472 (0.13, 4) | 46.8 | 86.2 |
| All layers | 666,444 (0.03, 3) | 100 | 767,466 (0.04, 4) | 100 | 86.8 |

### Neuronal number per areas MEC and LEC

A few estimates (n=2-6) were found reporting the number of neurons in EC areas MEC and LEC (**Suppl. Tables 11-14**). The results showed higher number of neurons in males (10-18% more) although a reversed trend was found in LEC, that had only one male datapoint and thus might not be reliable (**Table 9 and Figure 8B**). When analyzing the distribution of neurons across areas MEC and LEC, a quite consistent pattern is found in both sexes, with MEC having about 60% and LEC 40% of the neurons (**Table 10**). These percentages are further supported by additional data from (Heggland et al., 2015) (**Suppl. Table 1**) showing a very similar MEC/LEC ratio for mixed males and females (**Table 10**). In this regard, MEC and LEC estimates from (Rapp et al., 2002) suggest they used different segregation of MEC and LEC areas and were not used in the present analysis (**Suppl. table 1**).

**Table 9.** Number of neurons in MEC, LEC and EC by sex (from suppl Table 13). Males exhibited 10-18% more neurons than females in MEC and the whole EC, with a reversed trend in LEC, that had only one male datapoint. The average of males and females is the average of their means. Note: total number of neurons in Layers/areas/EC might vary across tables due to the use of different sets of data to obtain total averages

|  | MEC | LEC | EC |
| --- | --- | --- | --- |
| Average females (CV, n) | 384,099 (0.03, 3) | 283,548 (0.06, 3) | 667,647 (0.01, 3) |
| Average males (CV, n) | 467,195 (0.02, 2) | 264,800 (1) | 724,879 (0.10, 6) |
| Average M/F | 404,873 | 274,174 | 696,263 |
| Female/male (%) | 82.2 | 107.1 | 90.2 |

**Table 10.** Neuronal distribution across entorhinal areas. Data showed quite consistent distribution of neurons in MEC and LEC in both sexes, with MEC having about 60% and LEC another 40%. # Heggland et al., 2015 estimates were not used in the present analysis (Suppl. Table 1) but their data on mixed males and females is useful to obtain data the MEC/LEC ratio.

| % of total EC | MEC | LEC | EC |
| --- | --- | --- | --- |
| Females | 57.5 | 42.5 | 100 |
| Males | 64.2 | 35.8 | 100 |
| Average M-F | 58.1 | 39.4 | 100 |
| M-F (Heggland et al., 2015)# | 59.3 | 40.7 | 100 |

### Neuronal number per layers and areas MEC and LEC

Complete data describing the number of neurons in the different layers of both MEC and LEC was found only in 2 studies on females, that showed very consistent results (**Table 11 and Figure 14**), while we found only one study in males (Amaral et al., 2024) providing data for upper layers II/III and deep layers V/VI in MEC and LEC (**Suppl. tables 11-13**). From that combined layer II/III estimates we used the percentages reported in females (**Table 8)** to estimate the number of neurons in layer II (28%) and layer III (72%) that are shown in **Table 12 and Figure 9**. In addition, (Kumar and Buckmaster, 2006), reported number of neurons in males across layers but restricted to MEC (**Table 11 and Suppl. table 11-13**).

**Figure 9.**
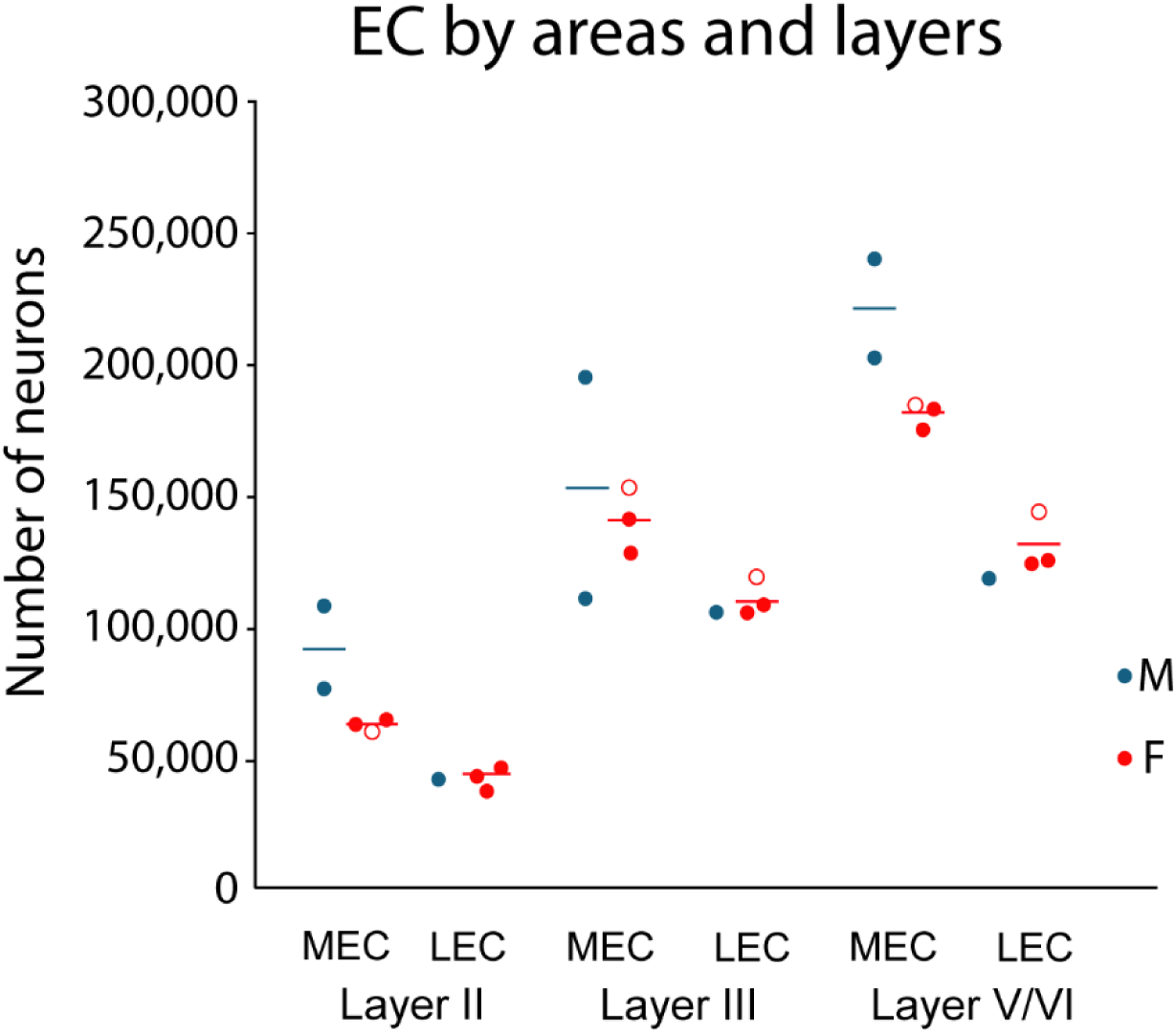
Number of neurons per area and layer in the EC for both sexes. Circles represent adolescent animals while circles represent adults. Averages are represented by horizontal lines. Note the striking consistency of female estimates.

**Table 11.** Number of neurons per layers and areas in the EC of females (from suppl tables 10, 11 and 12). Both the percentage of neurons per layer across areas, and the percentage of neurons in each area across layers is quite consistent.

| Females |  |  |  | % of all layers |  |  | % of all areas |  |  |
| --- | --- | --- | --- | --- | --- | --- | --- | --- | --- |
| Layer | MEC | LEC | EC | MEC | LEC | EC | MEC | LEC | EC |
| II | 61,719 | 41,635 | 102,150 | 16 | 15 | 15 | 60 | 40 | 100 |
| III | 140,801 | 110,796 | 251,597 | 37 | 39 | 38 | 56 | 44 | 100 |
| V/VI | 181,579 | 131,118 | 312,697 | 47 | 46 | 47 | 58 | 42 | 100 |
| total | 384,099 | 283,549 | 666,444 | 100 | 100 | 100 | 58 | 42 | 100 |

**Table 12.**
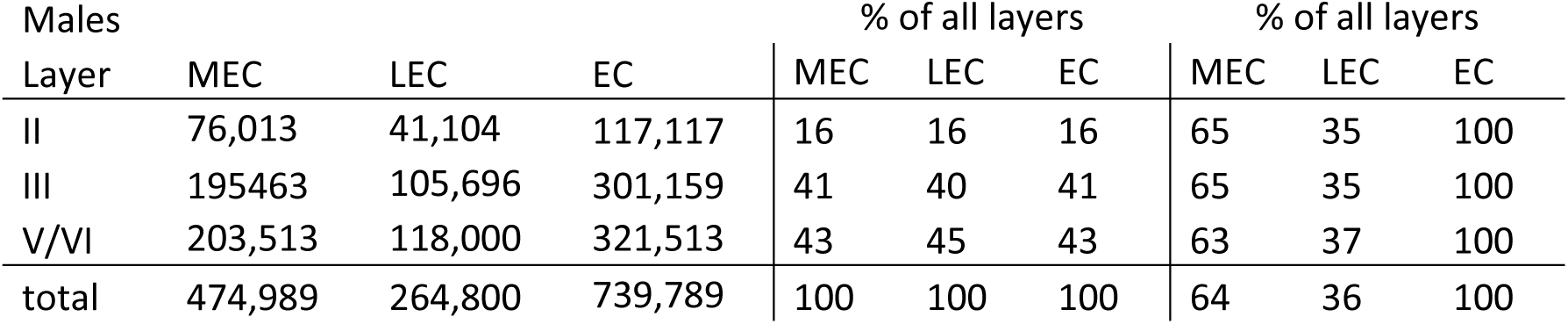
Number of neurons per layers and areas in the EC of males from Amaral et al., 2024 (suppl tables 10, 11 and 12). Both the percentage of neurons per layer across areas, and the percentage of neurons in each area across layers is quite consistent.

| Males |  |  |  | % of all layers |  |  | % of all layers |  |  |
| --- | --- | --- | --- | --- | --- | --- | --- | --- | --- |
| Layer | MEC | LEC | EC | MEC | LEC | EC | MEC | LEC | EC |
| II | 76,013 | 41,104 | 117,117 | 16 | 16 | 16 | 65 | 35 | 100 |
| III | 195463 | 105,696 | 301,159 | 41 | 40 | 41 | 65 | 35 | 100 |
| V/VI | 203,513 | 118,000 | 321,513 | 43 | 45 | 43 | 63 | 37 | 100 |
| total | 474,989 | 264,800 | 739,789 | 100 | 100 | 100 | 64 | 36 | 100 |

Analysis of the distribution of neurons per layers or areas showed consistent patterns, with layer II, III and V/VI exhibiting about 15%, 38% and 47% of the total in females, meaning upper layers represent 53% and deep layers 47% of the neurons. In males, similar rations were found, with upper layers showing 57% and deep layers 43% of the neurons. Analyzing neuronal population by area, both sexes showed similar ratios across layers, with females showing about 58% in MEC and 42% in LEC, and males having about 64% in MEC and 36% in LEC, suggesting an overall 60/40 split of the EC neurons between MEC and LEC. In this regard, an additional study (Heggland et al., 2015) using mixed males and females not included in the analysis (**Suppl. Table 1, Table 13**) showed that MEC had 59% and LEC 41% of the EC neurons, further supporting the 60/40 split of neurons between MEC and LEC.

Comparisons of the estimates by layer and area between males and females is not reliable given the low number of estimates involved. (**Table 12)**. In general, it suggests that males exhibit more neurons than females and the reverse trend in LEC might be an epiphenomenon of the only study available (Amaral et al., 2024) (see also **Table 9**).

Overall, the data from the entorhinal cortex shows surprising consistency across studies, particularly the data from females, even when originated from animals of different strain and age, including adolescents. There seems to be sex differences in the EC, with males having 10-15% more neurons than females (**Table 13**). Beyond that, both sexes show quite consistent distribution of neurons both across layers and between areas MEC and LEC. The EC has about 700,000 neurons, distributed 60/40 between MEC and LEC, and with approximately 15% in layer II, 38% in layer III and 47% in layers V/VI, proportions that were found to be virtually identical across sexes.

### Summary of the data

The estimated average of the number of neurons in each field of the hippocampal formation is provided in **Table 14**. Estimates for males and females are the average of all strains and are rounded for clarity and simplicity. Reliable estimates for females are provided for the GCL and EC, while estimates in parenthesis represent probable averages based on few datapoints including animals 1 month or older.

Regarding differences by strains, a summary of the averages of Wistar and SD adult males is provided in **Table 15** except for EC, that did not have enough datapoints for these strains. Wistar tends to show more neurons than SD, reaching statistical significance in CA3/CA2, although those differences are not consistent across fields and might result from small samples (subiculum) or potentially unbalanced averages (CA3/CA2). The most reliable data regarding number of datapoints and balanced samples may come from GCL and CA1, showing 5-10% differences that however do not reach statistical significance.

**Table 13.** Sex differences in the neuronal composition of the EC per layer and area, expressed as female percentage of the male value. The scarce number of studies available do not allow for reliable comparisons, but overall, the results suggest that males exhibit more neurons than females, and probably the inverse difference in LEC is due to the single value available for males.

| Layer | MEC | LEC | EC |
| --- | --- | --- | --- |
| II | 81.2 | 101.3 | 87.2 |
| III | 72.0 | 104.8 | 83.5 |
| V/VI | 89.2 | 111.1 | 97.3 |
| total | 80.9 | 107.1 | 90.1 |

**Table 14.** Summary of the data for all fields. Averages for males and females and 95% confidence intervals (CI). Data in italics indicates low reliability: CA3 estimate is based on only 4 datapoints; CA2 data are estimates (see text); MEC and LEC data from males come from one study and layer II and III data are estimates (see text) and thus not CIs are shown. Averages are rounded for simplicity and clarity and might not correspond to the midpoint of CIs.

|  |  | Adult males |  | Adult females |  |
| --- | --- | --- | --- | --- | --- |
|  |  | Average | 95% CI | Average | 95% CI |
| DG | GCL | 1,050,000 | 980,000-1,110,000 | 1.010,000 | 740,000-1,290,000 |
|  | hilus | 50,000 | 45,000-51,000 | - | - |
| CA | CA3 | ~210,000 | 163,000-268,000 | - | - |
|  | CA2 | ~30,000 | ~20,000-40,000 | - | - |
|  | CA3-CA2 | 240,000 | 214,000-260,000 | - | - |
|  | CA1 | 350,000 | 322,000-374,000 | - | - |
| SUB | Sub | 300,000 | 239,000-319,000 | - | - |
| MEC | II | 75,000 | - | 60,000 | 56,000-68,000 |
|  | III | 200,000 | - | 140,000 | 110,000-172,000 |
|  | V/VI | 200,000 | - | 180,000 | 169,000-194,000 |
| LEC | II | 40,000 | - | 40,000 | 29,000-54,000 |
|  | III | 100,000 | - | 110,000 | 93,000-129,000 |
|  | V/VI | 120,000 | - | 130,000 | 103,000-159,000 |

**Table 15.** Summary of the differences in the number of neurons between adult Wistar and SD males across fields. Differences are calculated as (1-(SD/Wi))*100 and statistical comparisons were performed with two-tailed Welch’s t-test. The average of SD animals in the subiculum is highlighted in italics as originates from only 4 estimates, producing a likely unreliable average and precluding statistical comparisons.

|  | SD | CV | n | Wistar | CV | n | Diff | Sign. |
| --- | --- | --- | --- | --- | --- | --- | --- | --- |
| GCL | 990,000 | 0.17 | 14 | 1,090,000 | 0.22 | 19 | 9% | p=0.16 |
| hilus | 47,626 | 0.21 | 15 | 48,097 | 0.11 | 13 | 1% | - |
| CA3/CA2 | 197,060 | 0.22 | 7 | 257,389 | 0.21 | 15 | 23% | p=0.014 |
| CA1 | 343,549 | 0.26 | 18 | 348,094 | 0.21 | 17 | 1.3% | p=0.87 |
| Sub | *242,133* | *0.23* | *4* | 308,466 | 0.09 | 5 | 22% | - |

For a summary and comparisons of all data by field, sex, strain and age (adolescent and adults) see Suppl. Table 15.

## Discussion

The hippocampal formation is essential for learning and memory, and its characteristic architecture, with distinct fields interconnected through unidirectional, spatially segregated projections, makes it an optimal candidate for models relating structure and function. A starting point of those models is the quantitative definition of the neuronal populations involved. The seminal report of (Amaral et al., 1990) provided one of the first comprehensive accounts of neuronal populations in the hippocampal formation and their connectivity, and quickly became the reference for most subsequent descriptions or models of the hippocampal formation, e.g.: (Treves and Rolls, 1992, 1994; Patton and McNaughton, 1995; Henze et al., 2002; Aimone et al., 2006, 2009; Leutgeb et al., 2007; Kempermann, 2011; Krueppel et al., 2011; Snyder and Cameron, 2012; Rolls, 2013; Newman and Hasselmo, 2014; Cole et al., 2020; Berdugo-Vega et al., 2023; Borzello et al., 2023; Vandael and Jonas, 2024). Since the data in that publication were based on the few available studies at the time, we have aimed to update those estimates with a comprehensive search for published studies. We reason that an update is useful to build more precise models of connectivity required for more precise inferences on function. We selected 87 studies reporting 264 estimates of principal neuronal populations in the main hippocampal fields, subiculum and upper and lower layers of the EC medial and lateral areas, plus some additional estimates of other subicular complex fields like pre- para- and post-subiculum. This curated dataset represents the most complete collection to date and provides updated estimates of hippocampal neuronal populations useful for modelling or establishing interspecies comparisons.

### Updated vs traditional averages

As mentioned above, most models and descriptions of the hippocampal formation (Treves and Rolls, 1992, 1994; Henze et al., 2002; Aimone et al., 2006, 2009; Leutgeb et al., 2007; Kempermann, 2011; Krueppel et al., 2011; Rolls, 2013; Newman and Hasselmo, 2014; Berdugo-Vega et al., 2023; Borzello et al., 2023; Vandael and Jonas, 2024) have adopted the populations described by (Amaral et al., 1990) for the tri-synaptic pathway: 200,000 neurons in EC layer II; 1,000,000 neurons in the GCL (Boss et al., 1985); 330,000 in CA3/CA2 (Boss et al., 1987) and 420,000 for CA1 (Boss et al., 1987). No data were available at the time for the EC, so their estimate of 200,000 neurons for EC layer II is a rough calculation based on the product of the areal density of neurons in EC layer II and the surface area of the EC, originally reported with a question mark to emphasize the approximate value of that estimate (see fig 1 in (Amaral et al., 1990)). They also described 30,000 in the hilus (Seress, 1988), about 12,000 neurons in CA2 and 128,000 in the subiculum based on (Cassell, 1980), but those estimates have not been used in subsequent models.

The results of our meta-analysis show an updated average for the GCL that is virtually identical to the traditional value of 1 million neurons. However, there are marked differences in the other populations. Our average of EC layer II describes half the number of neurons, while our updated average of CA3/CA2 and CA1 are 27% and 17% lower than the averages reported by (Boss et al., 1987), that are clearly outside the 95% confidence interval we obtained for both fields. While the difference in EC layer II is reasonable considering the rough methodology used to obtain the original estimate, the differences in CA fields are harder to explain, as the original estimates from (Boss et al., 1987) were obtained in 1 month-old SD females, and several studies have shown that adolescent age, female sex and SD strain typically correlate with lower number of neurons compared to male sex and Wistar strain, as described throughout the results. Furthermore, (Boss et al., 1987) described much larger neuronal populations in CA3/CA2 and CA1 (30-55% larger) in SD than in Wistar females, while our data suggest that Wistar might have significantly more neurons in CA3/CA2 and to a lesser extent, in other fields. We could not find an explanation for those discrepancies, but their study was not included in our sample, as they used assumption-based stereology.

In conclusion, it seems that the data from adolescent SD females from (Boss et al., 1987) might not be the most representative of the population of neurons in CA fields of the adult rat, and can be updated to more balanced estimates of 240,000 neurons for CA3/CA2 and 350,000 for CA1, based on the average of 24 and 37 estimates, respectively. Also, our analysis could not identify reliable studies describing the population of CA2, but we estimated, based on the relative volume and neuronal density of CA2 compared to CA3 and CA1, that CA2 might exhibit ∼30,000 neurons, implying that CA3 might have ∼210,000 neurons, in good agreement with the 215,000 neurons obtained from the few data available for CA3 alone.

While most models have relied in the data described above reported by (Amaral et al., 1990), there are others that have used partially different sources:

A model of the connectivity of the DG (Patton and McNaughton, 1995) used the 200,000 neuronal estimate for EC layer II and 1,000,000 for the GCL based on (Amaral et al., 1990), but updated the hilar number to 50,000 neurons based on (West et al., 1991), a value that matches the one we found in our meta-analysis.

Another model by (O’Reilly and McClelland, 1994) also used the estimate of 200,000 neurons for EC layer II and 1,000,000 for GCL, although they listed 850,000 GCs for their actual model. They described CA3 as having 160,000 neurons (Squire et al., 1989) likely after (Cassell, 1980), and CA1 as having 250,000 neurons, citing (Boss et al., 1987), although they probably meant (West et al., 1991). In addition to the difference in EC layer II already discussed, their values of CA3/CA2 and CA1 would represent 35% and 27% less neurons that the ones obtained in our analysis, and much lower percentages compared with the estimates of (Boss et al., 1987) commonly used in most models.

An additional model from the Snyder lab (Snyder and Cameron, 2012; Cole et al., 2020), described 112,000 and 247,000 neurons for EC layer II and III respectively after (Mulders et al., 1997); 1,200,000 neurons for the GCL and 250,000 for CA3/CA2, based on (West et al., 1991). This study is much closer to our results as the value for EC layers and CA3/CA2 are very similar to our estimates, while their GCL population is about 20% higher than the 1 million typically reported that we also found in our analysis.

In summary, models of the hippocampal formation have used neuronal populations that are typically based on single studies for each field. That implies that subjective factors in the choice of reference studies might largely determine the properties of the hippocampal network, and might produce vastly different results between models, as the quantitative relations between fields change dramatically. For example, it is hard to compare a model with 1 million GCs, 160,000 CA3/CA2 neurons and 250,000 CA1 pyramidal cells with another with the same number of GCs but double number of CA3 neurons (330,000) and almost 70% more CA1 neurons (420,000). Thus, to build more robust and accurate models, it seems reasonable to standardize neuronal populations, and the average of methodologically sound studies after removing outliers, seems the most reasonable option.

A recent meta-analysis by (Attili et al., 2022) collected a wide array of studies including stereological, physiological and neurochemical data to characterize quantitatively the different neuronal populations in the rat hippocampus. Their estimates of principal neuron populations are reported in their Figure 1D and in hipocampome.org: 962,000 neurons in GCL, 228,000 in CA3, 19,000 in CA2, 390,000 in CA1, 132,000 in Subiculum, 270,000 in LEC and 80,000 in MEC. Overall, some of their averages are within ∼10% of ours, but in others, the differences are remarkable, like in the subiculum (132,000 vs 300,000) or MEC (80,000 vs 265,000-285,000 depending on sex). A likely explanation for those divergences is that Attili and coworkers prioritize including as many studies as possible, without excluding outliers or filtering their source data for potential methodological issues; they pool data from animals of different sex, strain and age (e.g. mixing infants and adults) and also estimate neuron numbers from the product of neuron densities and volumes obtained in different animals/studies or incorporate mouse data to the model using a conversion factor obtained from the overall cortical neuronal counts in rat and mouse from (Herculano-Houzel et al., 2006). Thus, although an optimization algorithm for minimization of residuals is applied, it is possible that some of their estimates might not be accurate, as described above.

In this regard, high throughput models such as the one produced by Attili and coworkers or (Erö et al., 2018) in the mouse are useful tools to generate large datasets. However, by definition, models need to be compared with high quality datasets (when available) to evaluate their accuracy. And we think our analysis provides such comprehensive and likely accurate dataset on principal neuron populations in the rat hippocampus.

### Sex Differences

Studies on the hippocampus have traditionally favored the use of males to avoid the influence of cycling hormonal changes present in females, as the hippocampus is very responsive to hormonal fluctuations, sometimes in a specific manner (Gould et al., 1990; Zhang et al., 2008a). Indeed, this might be the reason behind the sexual dimorphism described in this structure: the male hippocampus is larger, heavier and contains more neurons than the female hippocampus (Madeira et al., 1992; Andrade et al., 2000; Nuñez et al., 2003b). The number of GCs has been reported to be higher in males in several strains of rats (Madeira et al., 1988, 1991; Nuñez et al., 2003a, 2003b; Schmitz et al., 2005; Smith et al., 2008) and studies in mice have shown that both sexes exhibit similar initial population of GCs early postnatally, that start differentiating during early adolescence (P20-P27) by increased cell death in females (Wimer et al., 1988). Sex dimorphism in the hippocampal formation is in agreement with the dimorphism in brain and body weight described in rats (Kakolewski et al., 1968; Goodrick, 1980; Madeira et al., 1988, 1991; Nuñez et al., 2003a, 2003b), mice (Wimer et al., 1988; Wimer and Wimer, 1989) and in guinea pigs (Severi et al., 2005), indicating this might be a general feature in rodents, and maybe in other species.

In our dataset, we did not find significant differences between sexes in the GCL, with males exhibiting similar neuronal numbers than females (within 5%). However, we found only 6 estimates from females, most using less common strains such as Long Evans and Fisher 344 and therefore the female average might not be reliable. As reported above, sex differences in the number of neurons have been extensively reported, and thus it seems a likely possibility although it was undetectable in our dataset. Additional studies reporting female data using robust stereological methods might be necessary to provide a more definitive answer.

Regarding other fields, several studies have reported males having 17-24% more neurons than females in CA1 and subiculum across different ages, from P10 to 6 months of age in both SD and Wistar animals (Madeira et al., 1992; Andrade et al., 2000; Smith et al., 2008), and differences in CA3/CA2 in 3 week-old rats (Nuñez et al., 2003a, 2003b). Also, (Smith et al., 2008) reported that SD females showed 7% more neurons than males at P9, but this difference disappeared in young adults (P68), consistent with data from (Madeira et al., 1992) describing similar averages in adolescent and adult SD males and females, suggesting sex differences in CA3/CA2 might subside in adulthood. In this line, (Schmitz et al., 2005) reported only slightly higher (4%) number of neurons in CA fields (CA1-CA3) of males compared to female rats 30-month-old. Our data lacks enough number of female datapoints to obtain reliable estimates, and therefore we could not make comparisons by sex. In the EC however, even when data were scarce for both sexes (n=3-6) we frequently found that males exhibited 10-15% more neurons than females, suggesting there might be indeed a sex difference that could be detected given the remarkable consistency of the data across studies in that field.

Given the potential for sex differences in most fields, we focused on obtaining data from males, the sex of choice of most studies on the hippocampal formation. However, when describing EC neuronal populations segregated by area and layer we focus on female data, that are limited but surprisingly homogeneous across studies, conferring them some seal of reliability, although such convergence might be just serendipitous given the small number of datapoints (n=2-5).

### Strain differences

The studies collected here used animals from 4 strains, but most studies used Wistar or Sprague-Dawley rats, so we focused the comparisons on those strains, and were limited to males, given the scarce data on females. The results show that Wistar typically exhibits more neurons than Sprague-Dawley males, although the differences are very negligible in some cases (within 5%) and inconsistent across fields.

The largest difference was found in CA3/CA2 (23%), while much more moderate 9% was found in the GCL and no difference was found in the hilus or CA1, while no reliable comparisons were possible in the rest of the fields. A closer view to the difference in CA3/CA2 reveals there are only 7 datapoints for SD animals. Considering the large variability of stereological estimates, this number of datapoints might provide a reliable average, but is also small enough to be influenced by particularly high or low estimates that do not qualify as outliers. In addition, the mild differences between strains found in other fields suggest caution when interpreting this major difference, that might be inflated by 2 particularly low values in the SD dataset. Thus, our analysis does not allow obtaining reliable conclusions, but it seems that the difference between Wistar and SD might not be substantial.

In contrast, some studies focused in the comparison between strains have shown marked differences. (Seress, 1988) analyzed LE versus CFY (a variety of SD) rats and showed that SD had ∼30% more neurons in GCL, but ∼7-10% less in hilus, CA2-CA3 and CA1, suggesting that potential differences across strains might not be consistent across fields, as our data show. However, the groups analyzed by (Seress, 1988) contained mixed sexes and therefore the strain effect might be confounded with sex differences as described in the previous section. (Boss et al., 1985, 1987) compared Wistar and SD females and found major differences between them. Their comparison in the GCL at 1, 4 and 12 months of age, showed that SD females had 45% and 28% more granule cells than Wistar at 1 and 12 months of age, but similar numbers at 4 months, while in CA3/CA2 Wistar had ∼36% less neurons (p<0.05) and 24% less in CA1 (p>0.05). These differences are substantial and show higher neuron number in SD than in Wistar, an inverse trend to that in our dataset. Comparisons within the same study are expected to be more reliable, as the same methodology is applied similarly by the same researchers, even if they used suboptimal assumption-based stereology. However, a point of contention is the results in the GCL, that are inconsistent across ages. Thus, no clear conclusions can be raised from those data.

### Age differences

Exploring neuronal numbers across age suggests two main questions: first, at what age hippocampal fields reach their adult complement of neurons, and second, if neuron numbers change with aging. To answer these questions, separate analysis must be made for the GCL and the rest of hippocampal fields, given their different neurogenic dynamics.

About the first question regarding when hippocampal fields reach their adult complement of neurons, the GCL shows protracted neurogenesis that reaches a peak at the end of the first postnatal week (Schlessinger et al., 1975; Bayer, 1980a) to subsequently decrease sharply during infancy and adolescence to reach low levels during adulthood (Arellano and Rakic, 2024). This continuous neurogenic environment complicates defining an adult number of neurons as a stable population across time. But modelling the process of neurogenesis reveals that at the onset of adulthood, by 2 months of age, neurogenesis generates ∼3.4% of the GCL population, a percentage that will decline following a power function to reach much lower levels afterwards (∼1% by age 6 months) (Arellano and Rakic, 2024), suggesting that the population will not change significantly after the onset of adulthood. Indeed, the same model and other studies have predicted that during the entire adulthood period between 2-21 months of age (Arellano et al., 2024), only ∼8-12% new neurons are produced in the rat (Ninkovic et al., 2007; Imayoshi et al., 2008; Lazic, 2012; Pilz et al., 2018; Arellano and Rakic, 2024). So, if there is no cell replacement, it would imply that during the adulthood period, the population of granule cells can be considered stable within a 12% margin, a relatively small difference that will go undetected considering the variability of the data across age, and is much lower than interindividual differences that can reach almost 2-fold, as discussed in the section “Variability, accuracy, and reliability of the data” below.

Comparisons in our dataset of the number of GCs in adolescent (0.75-2-months-old, (Arellano et al., 2024) and adults showed that adolescent males had about 8% less neurons than adults, although this estimated difference might not be reliable considering the limited number of adolescent datapoints (n=7). In this regard, available studies providing data on adolescent and adults show discrepant results, with Wistar male and females having about 20% less neurons than adults (Bayer et al., 1982; Boss et al., 1985), but adolescent Wistar males or SD females having the same or more than adults (Bayer et al., 1982; Heine et al., 2004), making difficult to interpret the overall trends.

Regarding the stability of the population of GCs across adulthood, our dataset does not allow a reliable analysis given the scarce data on middle aged and old animals and their large variability (∼1.6-fold variation between estimates at 12 and 24 months of age), that precludes regression (R2=0.05). An early study by (Bayer et al., 1982) quantified the number of GCs at 1, 4, 6.5 and 12 months of age, reporting a steady increase from 890,000 at 1 month to 1.28 at 12 months, implying a 30% increase or about ∼300,000 GCs, interpreted as the product of adult neurogenesis (Bayer et al., 1982; Crespo et al., 1986; Kempermann, 2011). However, with exception of Rasmussen et al., 1996, that reported a 20% increase between 2.5 and 24 months, all subsequent studies have not replicated those increases, showing stable or decreased numbers of neurons in older animals (Boss et al., 1985; Madeira et al., 1988, 1991; Rapp and Gallagher, 1996; Sousa et al., 1998; Merrill et al., 2003; Heine et al., 2004; Aslan et al., 2006). Also, available estimates of adult neurogenesis across adulthood (2-21 months (Arellano et al., 2024)) based on empirical data describe 8-12% of the population in the rat (Ninkovic et al., 2007; Imayoshi et al., 2008; Lazic, 2012; Pilz et al., 2018; Arellano and Rakic, 2024). Furthermore, as extensively discussed elsewhere (Arellano and Rakic, 2024), the addition of large numbers of neurons would imply the presence of large numbers of differentiating neurons expressing specific markers such as DCX or PSANCAM, in contrast with the modest numbers that have been described (Rao et al., 2005, 2006; Zhang et al., 2008b; Epp et al., 2009; Rennie et al., 2009).

Besides the GCL, the rest of hippocampal fields exhibit mostly embryonic neurogenesis that tappers during the first postnatal week (Bayer and Altman, 1974; Bayer, 1980a) producing a population peak around age 1 week, followed by neuronal pruning that leads to adult neuronal numbers by age 2 weeks (Bandeira et al., 2009). Hippocampal neurons differentiate along the first postnatal weeks and thus by the time animals reach adolescence at around 3 weeks of age (Arellano et al., 2024), it is expected that neurons occupy their adult positions and show adult-like soma morphology that would allow their quantification at similar rates than in adults. However, several studies quantifying the number of neurons in CA fields of adolescents (typically 1-month-old) and adult animals have shown consistently that younger animals exhibit less neurons (8-17% less) than adults, applying to both males and females (Madeira et al., 1992; Wakuda et al., 2008; Elibol-Can et al., 2014). A recent publication reporting number of neurons in CA1 and CA2 (combined) across development and adulthood showed a 5% difference between adolescents (1.25-month-old) and adults, and no difference between 1.4-month-old adolescents and adults suggesting no clear differences between both ages (De Araujo Furtado et al., 2024). Our dataset exhibited a trend showing lower neuron numbers in adolescents than in adults, but adolescent datapoints were very scarce (n<=5 per field), and in CA1, the field with more adolescent datapoints (n=5), we found, like (De Araujo Furtado et al., 2024), a mild 6% difference with adults that does not reach statistical significance (326,000 vs 247,000; p=0.061). Thus, there seem to be no clear explanation for the differences reported between adolescent and adult animals in CA fields, although those differences do not necessarily mean there are less neurons in adolescent animals but may indicate that a population of late migrating or late differentiating neurons might escape quantification in those fields, creating a mismatch between adolescent and adult neuronal numbers.

In contrast to the CA fields, the scarce data from the subiculum and EC was very homogeneous, without marked differences between adolescent and adults, or across sexes or strains (a feature that could be an artifact of the small number of samples). In the EC, given the small number of available estimates, we took advantage of that homogeneity and pooled one adolescent estimate with the adults to increase the total number of estimates.

The second question, regarding age-related loss of neurons in the hippocampal fields could not be answered in our dataset given the large variability of the estimates across ages and the scarce data from old animals (14% of datapoints). Thus, no regression could be found, precluding reliable temporal analysis. As discussed in each section, a number of studies have compared the number of neurons at different timepoints across adulthood (Andrade et al., 1995, 1996; Brandão et al., 1995; Rapp and Gallagher, 1996; Rasmussen et al., 1996; Merrill et al., 2003; Hattiangady et al., 2005; Heggland et al., 2015; Abhijit et al., 2018), and although some have reported sizable differences like 20% decrease in the hilus in old rats compared to young ones (Rasmussen et al., 1996) or a ∼30% decrease in CA1 between 6 and 18 months of age (Heggland et al., 2015), those differences were not statistically significant, given the large interindividual variation. Thus, it seems that hippocampal populations are essentially stable through adulthood.

### Rostro-caudal differences

Most studies report the overall number of neurons of the hippocampal fields, but the distribution of those neurons has been reported to change across the septo-temporal axis of the hippocampus. Traditionally, It has been considered that there are more GCs at septal levels than at temporal levels in adult Wistar males, and an inverse gradient is present for hilar neurons and CA3, with more neurons temporally than septally (Gaarskjaer, 1978; Buckmaster and Jongen-Rêlo, 1999; Amaral et al., 2007; Jiao and Nadler, 2007). However, (Wang and Gondré-Lewis, 2013) described similar number of neurons in CA3, but about double number of GCs and about triple number of CA1 pyramidal cells in the temporal half than in the septal half of the hippocampus in P14 mixed male and female SD rats. It is not clear if those different gradients might represent strain or age differences or might be due to methodological differences, but those mixed results suggest more studies are needed to clarify septo-temporal gradients.

### Variability, accuracy, and reliability of the data

The large variability between estimates observed in the current analysis is expected, as it is a typical feature of most stereological studies. There was a surprising exception in the EC, where a few estimates were particularly homogeneous, with coefficients of variation around 0.1, although this feature might be a serendipitous finding related to the small number of estimates in that region. As described along the manuscript, this variability precludes obtaining precise estimates or regression across age, and implies that, as indicated in the previous section, 20% or 30% differences between estimates from the same study will not be statistically significant.

The present analysis is a reviewed version of an initial analysis using a smaller and unfiltered sample of stereological studies (Arellano and Rakic, 2025). During the review process, it was suggested that more accurate and precise results could be achieved by curating the data to exclude those studies that used old, unreliable stereological methods (such as Abercrombie correction and other assumption-based designs) or had potential confounding factors such as counting nucleoli or performing unnecessary shrinkage corrections (**Suppl. Table 1**). After this initial filter, we realized that some of the selected estimates reported exceedingly large or small populations (e.g.: 1 million neurons in CA1) and those studies were also excluded (**Suppl. Table 1**). This was an interesting finding, as some of those results were obtained with modern, reliable stereological methods such as the optical fractionator, that would be expected to provide the most accurate results, but we could not identify potential sources of error.

At the end of the selection process, we had excluded almost half of the 166 studies initially identified, leaving 87 studies for analysis, a selected dataset that is expected to increase both accuracy and precision. Interestingly, there were only minor changes in the average number of neurons of each field, suggesting the original data was quite accurate. However, there was an improvement in the precision, as the confidence intervals of the average decreased in all fields subject to selection. Another consequence of the selection is that most of the data on females and on adolescent animals was excluded, as those studies were performed mostly using assumption-based stereology. Thus, the present study cannot make reliable comparisons between sexes or between adolescents and adults, as described in the next sections.

A final filter was applied to the selected data through the detection of outliers using Tukey fences (Tukey, 1977). Outlier detection might not be totally accurate when dealing with small samples as the ones described here. However, it seems an objective way to exclude values that are statistically too far from the average. The expectation is that, after eliminating outliers, the highest and lowest values will balance each other to produce a reliable estimate of the average population. However, given the small number of studies available, that balance is not warranted. An example is found when comparing strains in CA3/CA2, since adult SD males showed 23% less neurons than Wistar, a remarkable difference that reached statistical significance (p=0.014; two-tailed Welsch test). Although the difference seems reliable, based in 7 and 15 estimates respectively, adult Wistar males contain two high values (350,000, 360,000) coming from the (Rasmussen et al., 1996) study that also produced high estimates in CA1 (some of which are detected as outliers) while SD males contain low estimates coming from studies that also reported low values in other fields, like (Chen et al., 2008; Keleş et al., 2019). Thus, the major difference detected might be accentuated by those studies, and when removed, the difference is decreased to 12% and is not statistically significant. However, there are no objective reasons in the methodology or results of those studies to be excluded from the analysis, so we included them in the average. The main objective of this analysis is to obtain balanced averages that might compensate for high or low values produced in individual studies. This might not be achieved in every case, particularly when the number of datapoints is relatively low, as in this case (n=7), but still seems to be the best (or the least bad) approach to get the most balanced results. In that regard, we describe the average of all adult males, that increases substantially the number of estimates and produce a more balanced and accurate estimate for the overall population.

In addition to the average, we have calculated the median, a source of valuable information on the symmetry of the data after removing outliers. The median was similar to the average in every field (within 4%) except in the subiculum. In this field, most estimates orbited around 300,000 neurons, except in two studies that reported averages below 200,000 that lowered the average to 280,000. However, the median seemed a better estimator, reporting 296,000 neurons, and this estimate was therefore selected as the number of neurons in the subiculum.

We explored the possibility of weighing estimates according to the number of animals used to obtain them. This approach seems reasonable as averages based on more animals are expected to be more representative and have less variance. However, weighing the estimates did not produce significantly different averages (1-2% difference) and more importantly, we could not find any correlation between the number of animals used to obtain an estimate, and its proximity to the overall average. In other words, some of the studies with larger number of animals produce estimates that are far from the average and sometimes are suspected outliers, as occurs for example in the subiculum with two estimates below 200,000 neurons, about 50% below the 300,000 neurons reported by most other studies. Thus, the accuracy of the studies is not consistently related to the number of animals used, but to other factors related to the specific methodology and criteria used in each study. Therefore, we did not weigh the estimates, as it would not significantly change the average, but is another layer of complication and might produce small but unpredictable deviations of the average.

The large variability observed in the studies collected might originate from biological or methodological differences. Large biological differences are not expected from inbred animals of the same sex and strain and similar age, although minor differences between cohorts of animals obtained from different providers and bred in different laboratories under slightly different environmental conditions might produce some variability. However, we think that it is more likely that methodological differences across labs might exert a much larger contribution. In this regard, stereological quantification of neuronal populations using adequate methodology have reported ∼50% differences (480,000 vs 730,000) between estimates of GC neurons from adult male mice littermates, and almost 100% in 2-month-old littermates of both sexes (Ben Abdallah et al., 2010).

Those major differences are likely not related to actual biological differences, but to inherent imprecision of the stereological methodology, that nonetheless achieves a reasonable sample average within adequate statistical margins of error. Thus, if estimates from individual littermates analyzed in the same lab, with the same methodology and by the same researcher (L. Slomianka, personal communication) can have such large range of variation, it might not surprising that results can exhibit relatively large differences across labs given their methodological differences, using different stereological schemes, sampling methods and different criteria to divide hippocampal fields or to identify neurons to be counted, or to exclude non-neuronal cells might produce shifts of the averages up and down across studies. Thus, methodological factors might be the main contributor to the variability normally observed in stereological quantifications, although it does not seem possible to pinpoint specific factors driving those changes.

Optimally, estimates of number of neurons in each field should be obtained for each sex and strain separately. However, the scarce data for females and the relatively small number of estimates for the different strains produce less reliable results, as for example the significant difference reported in CA3/CA2 between SD and Wistar discussed above. Thus, we have focused the results on the male population, regardless of strain. Exceptionally, we estimated the population of the EC based on female data as we are interested in the number of neurons per area and layers, and the data available comes from females (although we estimated the populations for males based on one study).

The goal of this analysis is to obtain the most representative and hopefully, accurate estimates of neuronal populations in the hippocampus to help build updated and accurate connectivity models and provide reliable numbers of interspecies comparisons. Hopefully, the artificial intelligence revolution might bring automatization to stereological quantification that might allow mass production of quantitative studies, providing more definitive answers regarding neuronal populations and differences across strain and sex. Until then, we think the present report provides the most complete resource and analysis on the neuronal populations in the hippocampal formation of the rat.

## Conclusion

As part of a larger effort to review and update the model of connectivity of the hippocampal formation, we present here a comprehensive collection of studies reporting the number of neurons in the fields of the hippocampal formation. A relevant feature of the data is that is mostly restricted to adult males, with scarce data from females and adolescent animals, making difficult to compare data across age or sex. Sexual dimorphism has been reported in the hippocampus of rodents, with larger and more populated fields in males. The few estimates from the EC consistently suggest that males exhibit 10-15% more neurons than females, but no definitive conclusions can be raised. This difference might be present across hippocampal fields, but the scarce data on females precludes reliable comparisons. Another outstanding feature is the variability of the estimates across fields, with the exception of the EC, that also complicate comparisons across strains.

Comparison of our results with neuronal populations in existing hippocampal models shows marked differences in most fields that reach 2-fold differences, as existing models have obtained their data from individual studies that might not be representative. We propose that more reliable values can be obtained by averaging all data available, that produces the most representative and “least inaccurate” measure available. We hope these estimates will be useful to improve comparisons across species and to build more accurate models of hippocampal function.

